# Deep intravital brain tumor imaging enabled by tailored three-photon microscopy and analysis

**DOI:** 10.1101/2023.06.17.545350

**Authors:** Marc Cicero Schubert, Stella Judith Soyka, Amr Tamimi, Emanuel Maus, Robert Denninger, Niklas Wissmann, Ekin Reyhan, Svenja Kristin Tetzlaff, Carlo Beretta, Michael Drumm, Julian Schroers, Alicia Steffens, Jordain Walshon, Kathleen McCortney, Sabine Heiland, Anna Golebiewska, Felix Tobias Kurz, Wolfgang Wick, Frank Winkler, Anna Kreshuk, Thomas Kuner, Craig Horbinski, Robert Prevedel, Varun Venkataramani

**Author notes:** These authors contributed equally (Marc Cicero Schubert, Stella Judith Soyka, Amr Tamimi). These authors contributed equally (Robert Prevedel, Varun Venkataramani).

## Abstract

Intravital two-photon microscopy has emerged as a powerful technology to study brain tumor biology and its temporal dynamics, including invasion, proliferation and therapeutic resistance in the superficial layers of the mouse cortex. However, intravital microscopy of deeper cortical layers and especially the subcortical white matter, an important route of glioblastoma invasion and recurrence, has not yet been feasible due to low signal-to-noise ratios, missing spatiotemporal resolution and the inability to delineate myelinated axonal tracts. Here, we present a tailored intravital microscopy and artificial intelligence-based analysis methodology and workflow that enables routine deep imaging of glioblastoma over extended time periods, named Deep3P. We show that three-photon microscopy, adaptive optics, as well as customized deep learning-based denoising and machine learning segmentation together allow for deep brain intravital investigation of tumor biology up to 1.2 mm depth. Leveraging this approach, we find that perivascular invasion is a preferred invasion route into the corpus callosum as compared to intracortical glioblastoma invasion and uncover two vascular mechanisms of glioblastoma migration in the white matter. Furthermore, we can define an imaging biomarker of white matter disruption during early glioblastoma colonization. Taken together, Deep3P allows for an efficient and non-invasive investigation of brain tumor biology and its tumor microenvironment in unprecedented deep white and gray matter of the living mouse, opening up novel opportunities for studying the neuroscience of brain tumors and other model systems.

## Introduction

Glioblastomas (GB) are the most common malignant, primary brain tumors characterized by their infiltrative growth and colonization of the entire normal brain^1, 2^. Furthermore, these tumors are cellularly and molecularly heterogeneous^3–6^ with a notorious therapeutic resistance towards standard-of-care treatment with radio– and chemotherapy as well as surgical resection^7^. It has long been known that GB are predominantly a disease of the white matter^8^. GB frequently occur in the white matter and can invade into the contralateral hemisphere along the corpus callosum^9^. Furthermore, a majority of glioblastoma recurrence is detected within the white matter^9^. Recently, a large autopsy series further highlighted the importance of white matter tracts as an invasion route to invade and colonize the brainstem^1^. However, experimental studies of glioblastoma in the white matter have so far been entirely restricted to *ex vivo* analyses^10–12^ as technologies for deep intravital and longitudinal monitoring were lacking. This highlights the need and importance of studying this highly dynamic disease *in vivo* in the microenvironmental niche of the white matter. Such an approach would allow dissecting the underlying principles of glioblastoma biology including the yet elusive mechanisms of cellular invasion in the corpus callosum.

Intravital imaging of brain tumors including gliomas as well as brain metastases has so far been restricted to superficial cortex layers, dictated by two-photon microscopy (2PM) which is fundamentally limited to an effective penetration of ∼300-700 μm due to scattering and out-of– focus fluorescence background^3, 6, 13–17^. Deep *in vivo* brain imaging with 2PM in the mouse brain is currently only possible using highly invasive methods such as gradient index lens implantation or cortical aspiration. These methods have not yet been used to investigate intravital brain tumor biology because of the associated invasiveness.

In this respect, three-photon microscopy (3PM) has shown potential for deeper imaging beyond 1 mm in the brain due to an increased signal-to-background ratio and longer wavelength excitation which reduces tissue scattering^18, 19^. So far, 3PM has been used to investigate the static morphology of brain vasculature and neurons^18, 20^, and to perform calcium imaging of neurons^21^ and astrocytes^22^. Yet, their application to investigate spatiotemporally dynamic brain tumor biology poses additional technical challenges. In particular, tumor masses are opaque and dense structures, leading to a significant scattering of light^23^. Furthermore, the fluorescent membrane labelling of tumor cells required to visualize fine processes^3^ leads to intrinsically low fluorescence signals. This is exacerbated by the relatively high density of the labelled cells within a given imaging volume, which makes distinguishing and tracking of individual tumor cells and their fine cellular processes extremely difficult. With this comes the requirement of long-term stability of the microscopy system retaining high spatial resolution during extended volumetric time-lapse imaging and the ability to track the same image regions over days and weeks, all while retaining high spatial resolution. Lastly, photodamage– and toxicity need to be prevented by diligent optimization of laser power while keeping a reasonable signal-to-noise ratio (SNR), and an efficient screening system needs to be established that allows to identify both appropriate tumor regions and the white matter niche within a minimal time, in order to utilize imaging time and laser power diligently.

As outlined above, a predominant challenge in deep tissue imaging, including 3PM, is the low SNR of the raw image data. Here, computational methods using deep learning-based image restoration^24, 25^ have the potential to substantially increase the low SNR of images typically acquired in deep tissue conditions. While now increasingly applied to images obtained with confocal, light-sheet or two-photon microscopy setups^3, 25^, these methods have not yet been adapted and customized to the peculiar detector noise and background sources typically encountered in 3PM. Another opportunity of 3PM lies in the label-free imaging of blood vessels and myelinated axonal tracts in the brain using the third-harmonic generation signal. As these are both important structures of the tumor and brain microenvironment, it would be desirable to unequivocally demix and classify the THG signal into both anatomically distinct structures.

To overcome all of the above-mentioned fundamental technical limitations, we further adapted and advanced state-of-the-art 3PM and image analysis methods to enable the study of glioblastoma biology in the gray and white matter at unprecedented depths. In particular, we developed a minimally invasive intravital imaging and analysis workflow for modified patient-derived glioblastoma xenograft models stably transduced with a membrane-bound green fluorescent protein. Using bespoke 3PM, adaptive optics, and AI-enhanced post-processing and analysis tools, we demonstrate the capability to investigate glioblastoma biology and its tumor microenvironment in unprecedented deep white and gray matter of the living mouse, up to a depth of 1.2 mm and at near diffraction-limited spatial resolution. Deep3P time-lapse imaging allowed us to uncover distinct behavioral differences of glioblastoma in white matter tracts as compared to the gray matter of the cortex. Furthermore, we were able to track myelinated axonal fibers over weeks and characterize how they change in the course of early white matter glioblastoma colonization. In particular, Deep3P allowed us to compare invasion patterns in the whole cortex and subcortical white matter, which revealed an enrichment of a vascular invasion route from the cortex into the corpus callosum as compared to intracortical invasion. Within the corpus callosum, we found that glioblastoma cells and their neurite-like processes predominantly align with white matter tracts and follow the anatomical structure of myelinated axons. Surprisingly, we uncovered two additional vascular invasion mechanisms within the corpus callosum allowing invasion orthogonal to the fiber tracts similar to patterns from oligodendrocyte and astrocytic precursor cells during neurodevelopment^26, 27^. Lastly, the third harmonic generation (THG) signal of our Deep3P methodology permitted to dynamically investigate white matter disruptions during glioma infiltration of the corpus callosum. Here, morphometric characterization of the THG signal indicated an imaging biomarker of white matter disruption during early glioblastoma colonization which extends the applicability of our workflow to investigate axonal degeneration in other tumor and disease models.

## Results

### Infiltration of the corpus callosum as a hallmark of glioma

Previous research has suggested that infiltration into the white matter tracts may be an important route for glioblastoma to invade the contralateral cortex^12, 28^. However, it was not clear how prevalent the infiltration of the corpus callosum is in a human patient glioma cohort. To address this question, we analyzed a large brain tumor patient autopsy cohort (n = 50 patients) and found that 84% of all glioma patients showed at least microscopic infiltration into the corpus callosum irrespective of the location of their primary clinical manifestation (Fig. 1a, Extended Data Fig. 1a-e, Supplementary Table 1). Further analysis showed that 82% of patients with isocitrate dehydrogenase (IDH)-wild-type (n = 47 patients) and 100% of patients with an IDH-mutant glioma showed corpus callosum infiltration. In addition, we studied 20 patient-derived xenograft as well as patient-derived organoid xenograft glioma models and found that all of the analyzed models demonstrated a microscopically visible, infiltrative behavior of the corpus callosum after 40-60 days of implantation, regardless of whether the tumor was implanted into the cortex or striatum (Fig. 1a, Extended Data Fig. 1f, Supplementary Table 2). These findings indicate that white matter infiltration of the corpus callosum is a hallmark of glioblastoma growth, highlighting the need for intravital imaging of glioblastoma invasion and colonization inside the distinct microenvironment of the corpus callosum (Extended Data Fig. 1g), which is yet unattainable with current light microscopy technologies.

**Fig. 1.**
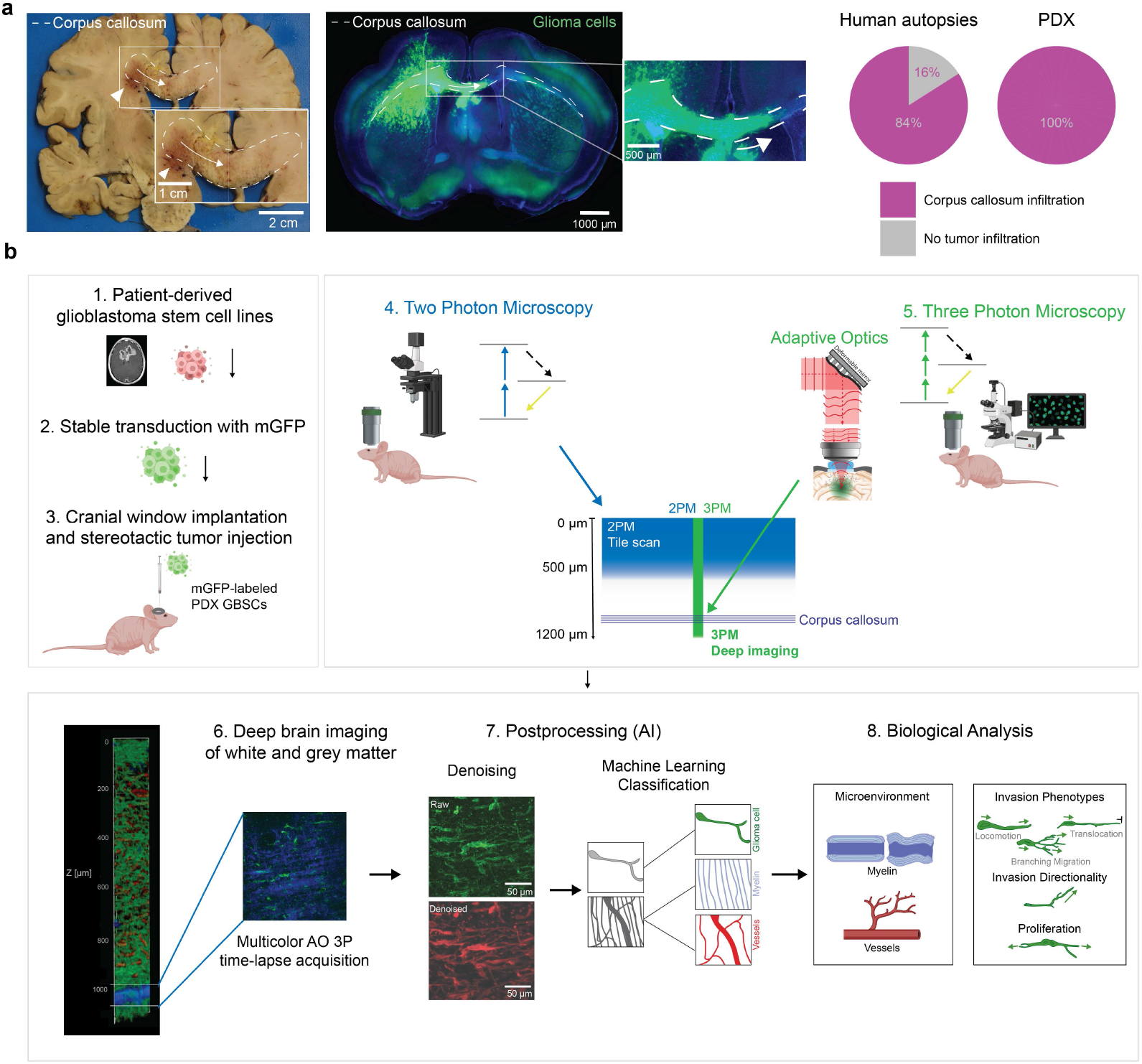
Rationale and combined experimental/analysis pipeline for deep brain tumor imaging. a, Left: Autopsy brain slice of a patient with IDH-WT glioblastoma. The corpus callosum is labeled with a dashed line. Arrowhead indicates main tumor mass; arrow indicates tumor infiltration in the corpus callosum as seen in the inset. Middle: Tumor infiltrated corpus callosum in S24 PDX mouse model. The corpus callosum is labeled with a dashed line, fluorescently labeled glioma cells are shown in green. Arrow indicates glioma infiltration of the corpus callosum. Right: Percentage of corpus callosum infiltration in human autopsies (n = 43 patients) and in PDX models (n = 23 PDX models). **b,** Workflow of Deep3P. Upper left: Establishment of *in vivo* patient-derived glioma xenograft model. Glioblastoma cell lines are derived from patients and stably transduced with mGFP and injected into mice brain. Upper right: Combination of 2PM and 3PM to identify target regions for deep brain imaging. Bottom: Deep brain imaging in the corpus callosum and subsequent deep and machine learning based post-processing to allow simultaneous myelin, vessel and tumor analysis.

### Deep3P as a novel workflow to investigate deep brain tumor biology

To investigate glioblastoma biology in the cortex and subcortical white matter tracts, we used state-of-the-art patient-derived xenograft models that accurately reflect glioblastoma molecular characteristics as analyzed with methylation array analyses and the histological growth patterns of glioma patients (Fig. 1a, Extended Data Fig. 1e, Supplementary Table 2). To visualize the fine processes of glioblastoma cells (GBCs) *in vivo*, we stably transduced them with lentivirally packaged membrane-bound GFP (mGFP) and injected them into the cortex of adult mice at a depth of approximately 500 µm (Fig. 1b). We used two-photon microscopy (2PM) to longitudinally screen the injected cells up to a depth of 600-700 µm in large diagonal field of views of up to 4.3 mm, taking advantage of the higher imaging speed of 2PM for mapping dynamic glioblastoma growth and its tumor microenvironment in superficial layers to select appropriate regions for deep imaging (Fig. 1b).

We then used our bespoke Deep3P methodology (Fig. 1b) to study the previously identified glioblastoma infiltration zone and investigate glioblastoma invasion, proliferation and colonization in the corpus callosum up to a depth of 1200 µm, a depth that is far outside the reach of 2PM (Fig. 2a). In particular, 3PM results in significantly enhanced signal-to-noise (SNR) ratios at all imaging depths (Fig. 2b). Deep3P leverages recent technological advances in 3PM and the addition of modal-based, indirect adaptive optics^22^ to optimize fluorescence signal and ensure near diffraction-limited performance at large imaging depths. The system is based on a custom-built 3P laser scanning microscope that was specifically optimized for 1,300 nm excitation and the use of short-pulsed (∼70 fs), low repetition-rate (<1 MHz) lasers that are critical to obtain large imaging depths^20, 22^. To improve the 3PM signal and imaging resolution, we employ a custom, modal-based indirect adaptive optics (AO) approach to correct for wavefront aberrations occurring due to the refractive index (RI) differences between the water-immersion, cranial window and brain, as well as brain tissue intrinsic RI inhomogeneities^22^. To successfully apply this approach to the requirements of intravital and longitudinal monitoring of cellular dynamics and white matter structural changes, the following challenges had to be met: (1) The entire imaging system and workflow were optimized for non-invasiveness, so that deep-brain time-lapse imaging over multiple hours did not induce photo-damage in terms of bleaching and/or toxicity. This also included adaptations to the cranial window surgery, mounting to minimize light loss due to reflections and absorption, as well as the establishment of an optimized imaging workflow (see Methods and Supplementary Note 1). (2) The fast AO correction measurement speed as well as robustness of the modal-based indirect AO approach^22^ was paramount to enable time-lapse volumetric recordings with little time overhead, while minimizing the overall light-exposure (Fig. 2c). Here, AO led to an average improvement in effective resolution of 1.9±0.2-fold (n = 5 image volumes, see Methods) and a 3-5-fold enhancement of fluorescence signals, as evidenced by intensity line plots and spectral power map analysis of lateral resolution (Fig. 2d-f). In effect, fine biological structures such as neurite-like processes of glioblastoma cells called tumor microtubes (TMs), blood vessels and cell nuclei can be resolved (Fig. 2g).

**Fig. 2.**
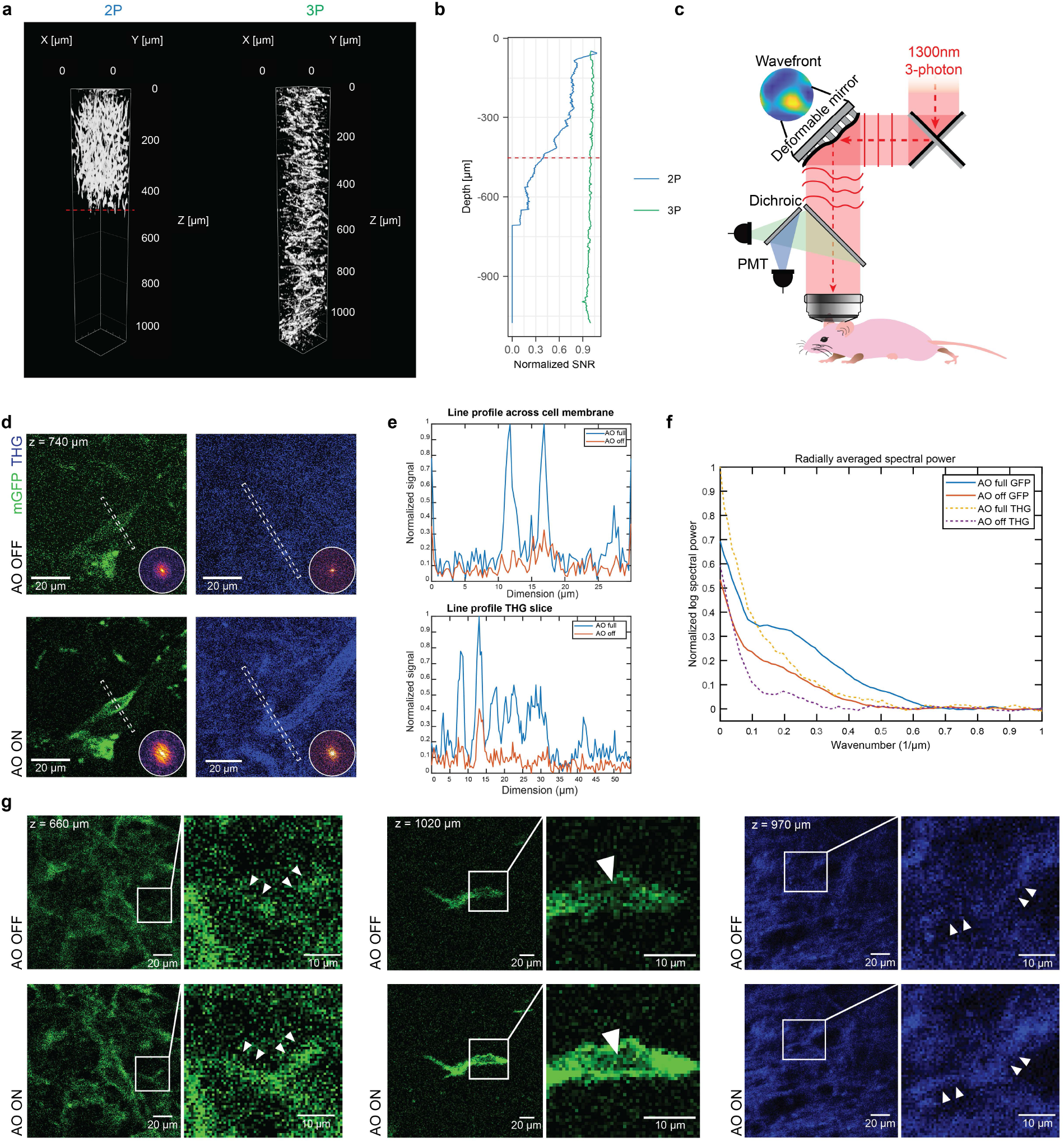
AO-enhanced 3PM and AI-based denoising allows near-diffraction limited resolution at large image depths and high SNR imaging. a, 3D renderings showing 2PM and 3PM imaging down to approximately 1000 µm. Gamma values were adjusted for 3D visualization. **b,** SNR of 2P and 3P along depths, normalized to the SNR at the brain surface. The dashed red line in **a-b** indicates the imaging depth at which biological structures cannot be clearly discerned anymore in 2PM in contrast to 3PM (approximately 450 µm). **c**, Scheme of 3PM and adaptive optics setup. **d**, An example of a glioma cell imaged without (top left) and with (bottom left) AO optimization and the corresponding images on the THG channel (top and bottom right). The line indicates the line segment averaged over to produce the line profiles in **e**, showing the effect of uncorrected optical aberration on the visibility of fine cellular structures. The inset on each panel shows the frequency domain power spectrum of the image, with the ring indicating 1 µm length scale. The image with adaptive optics shows significantly higher frequency contributions as compared to the image without adaptive optics. **e,** Line profile comparisons for both mGFP and THG channels showing intensity enhancement. **f,** The averaged radial profile of the frequency maps is shown, allowing easier estimation of the respective frequency cut-offs. **g,** Exemplary 3PM images with and without AO optimization on the mGFP channel at different depths in deep cortex and corpus callosum (left and middle) and on the THG channel within the corpus callosum (right). Zoom-ins are shown on the right. Arrowheads show TMs (left), a shadow of a cell nucleus that is caused by the membrane-bound GFP labeling (middle) and a blood vessel (right) on AO on and off images. TMs, the cell nucleus and the blood vessel are not clearly visible without AO.

### AI-based image denoising and analysis of Deep3P data

To further improve the signal-to-noise ratio of the raw images during time-lapse imaging of brain tumor cells while keeping 3PM excitation power minimal, we implemented a customized deep learning-based denoising workflow that can also account for the peculiar, structured noise often encountered in 3PM images (Fig. 3a-c, Extended Data Fig. 2).

**Fig. 3.**
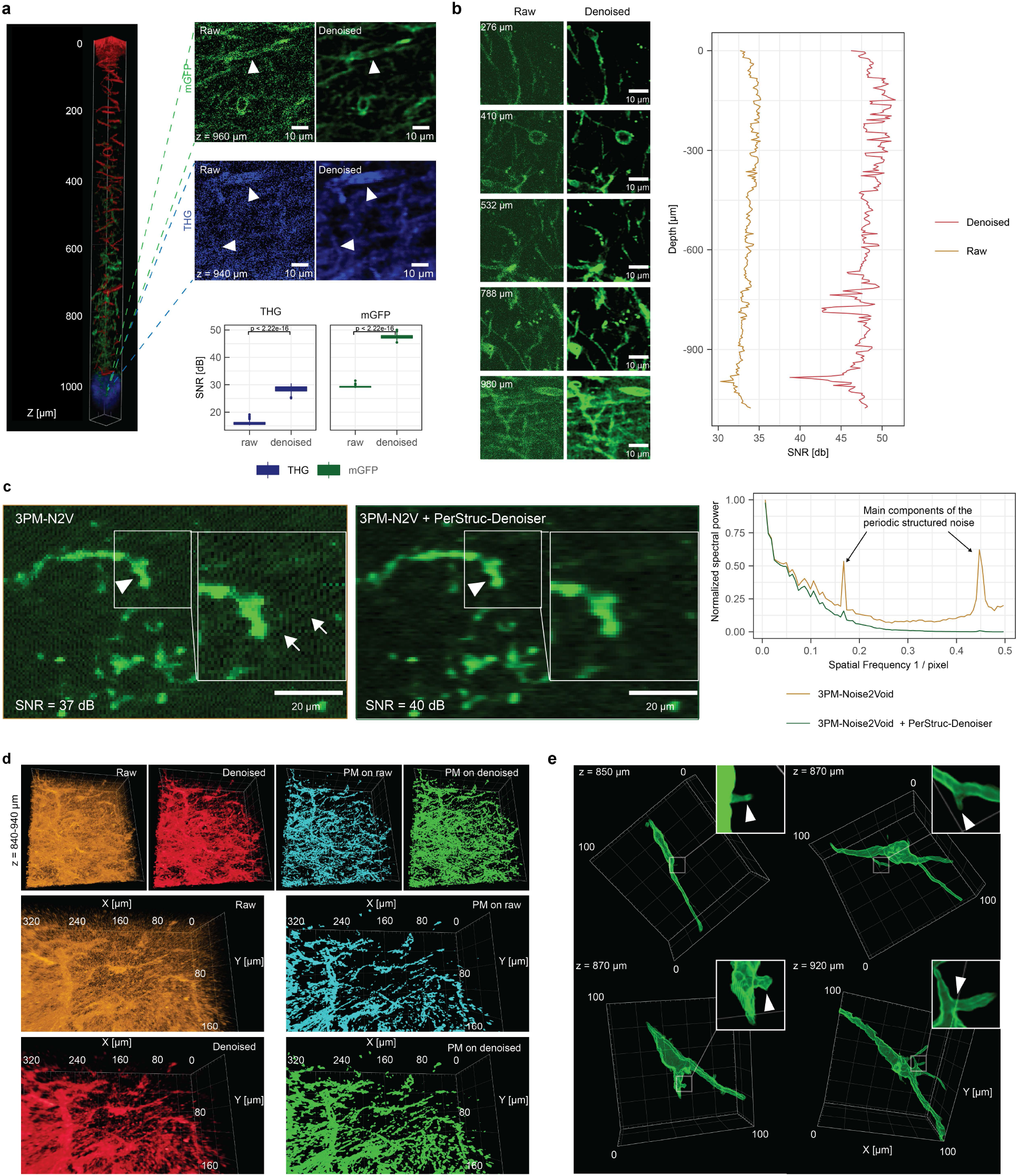
Denoising of AO-3PM and subsequent machine learning allows brain tumor imaging across the entire cortex and corpus callosum. a, Top left: 3D rendering of a stack going from the surface down to the corpus callosum. Red: blood vessels, blue: corpus callosum, green: GBM cells. Based on probability maps. Top right: Comparison of raw (left) and denoised (right) images within the corpus callosum (dashed lines on the left indicate imaging depth). Arrowheads in the mGFP image point to a glioblastoma cell soma that can be barely seen without denoising. Arrowheads in the THG signal point to fibrous structures that can be clearly identified after denoising. Bottom: signal-to-noise ratio (SNR) comparison of raw and denoised in THG and mGFP signal. Mann-Whitney-test. **b,** Left: Exemplary raw and denoised images of glioblastoma cells at different depths. Right: SNR in raw and denoised images across entire image stack. **c,** Comparison of the denoised 3PM-N2V image (left) and its version with additional application of the PerStruc-Denoiser (middle) showing the qualitative improvement corresponding to a 3 dB increase in SNR allowing a clearer identification of TMs (arrowhead). Arrows point to structured noise. Right: Averaged line power spectrum of the images depicting the PerStruc-Denoiser’s suppression of the main components of the periodic structured noise (see arrows pointing to its main components). **d,** 3D renderings based on raw images, denoised images, probability maps based on raw images and probability maps based on denoised images. **e,** Close-up 3D renderings of single glioma cells based on probability maps. The arrow heads on the zoom-ins point at small processes (top images and bottom left image) and a TM branching point (bottom right image). Gamma values were adjusted for 3D visualization in **a**, **d**, **e**.

In particular, we developed a bespoke 3D variant of the Noise2Void approach^24^ for the denoising of 3PM data, which utilizes a 3D U-Net architecture (Extended Data Fig. 2). Our method, referred to as 3PM-Noise2Void, effectively improves the image quality by taking into account the 3D nature of the data. This workflow increased the SNR by an additional ∼10-15 dB across all imaging depths (Fig. 3a-b, Supplementary Video 1). Furthermore, to counteract periodic structured noise patterns that cannot be addressed by Noise2Void, we implemented an additional post-processing technique, called PerStruc-Denoiser, to further reduce this structured noise effect by ∼3 dB (Fig. 3c, Extended Data Fig. 2). Lastly, we implemented a machine learning-based segmentation using the denoised data as a prediction mask for further biological analysis (Fig. 3d).

Furthermore, we highlight that denoising on 3PM images without AO resulted in qualitatively and quantitatively worse image quality compared to 3PM images with AO correction on (Extended Data Fig. 3). The effective increase in image SNR due to our bespoke AO and AI-based denoising is key for our Deep3P workflow. It allowed to keep excitation power and hence photodamage low during the extended, up to 4 hours long, time-lapse image acquisitions. In effect, this maintained and ensured the non-invasiveness of our imaging approach. Overall, the improvements in resolution and SNR of Deep3P across all imaging depths were critical to resolve the fine, neurite-like, cellular protrusions of the glioblastoma cells (Fig. 3e) and to qualitatively as well as quantitatively distinguish between different cellular migration and invasion patterns *in vivo*.

Our Deep3P imaging system permits acquisition of two separate channels, which were used for mGFP to visualize tumor cells, and to record the THG signal. The latter yields label-free image contrast that visualizes both myelinated white matter tracts, as well as blood vessels in the brain^29, 30^. Since white matter tracts and blood vessels constitute important tumor microenvironmental niches^8, 28^, we aimed at unambiguously differentiating the THG signal.

To this end, we trained a customized, interactive machine learning algorithm based on ilastik^31^ (see Methods) to distinguish between the background, blood vessels, and myelinated axonal tracts (Fig. 4). This enabled us to investigate the bidirectional relationship between brain tumor cells and the blood vessel and white matter microenvironmental compartments (Fig. 4a-d, Supplementary Video 1-3). We validated our AI-classifier by intravenous injection of FITC-dextran as ground truth signal for blood vessels (see Fig. 4e-g and Methods). This workflow allowed to clearly distinguish between blood vessels and myelinated axonal tracts both in 2D (Fig. 4c) and 3D rendering (Fig. 4d).

**Fig. 4.**
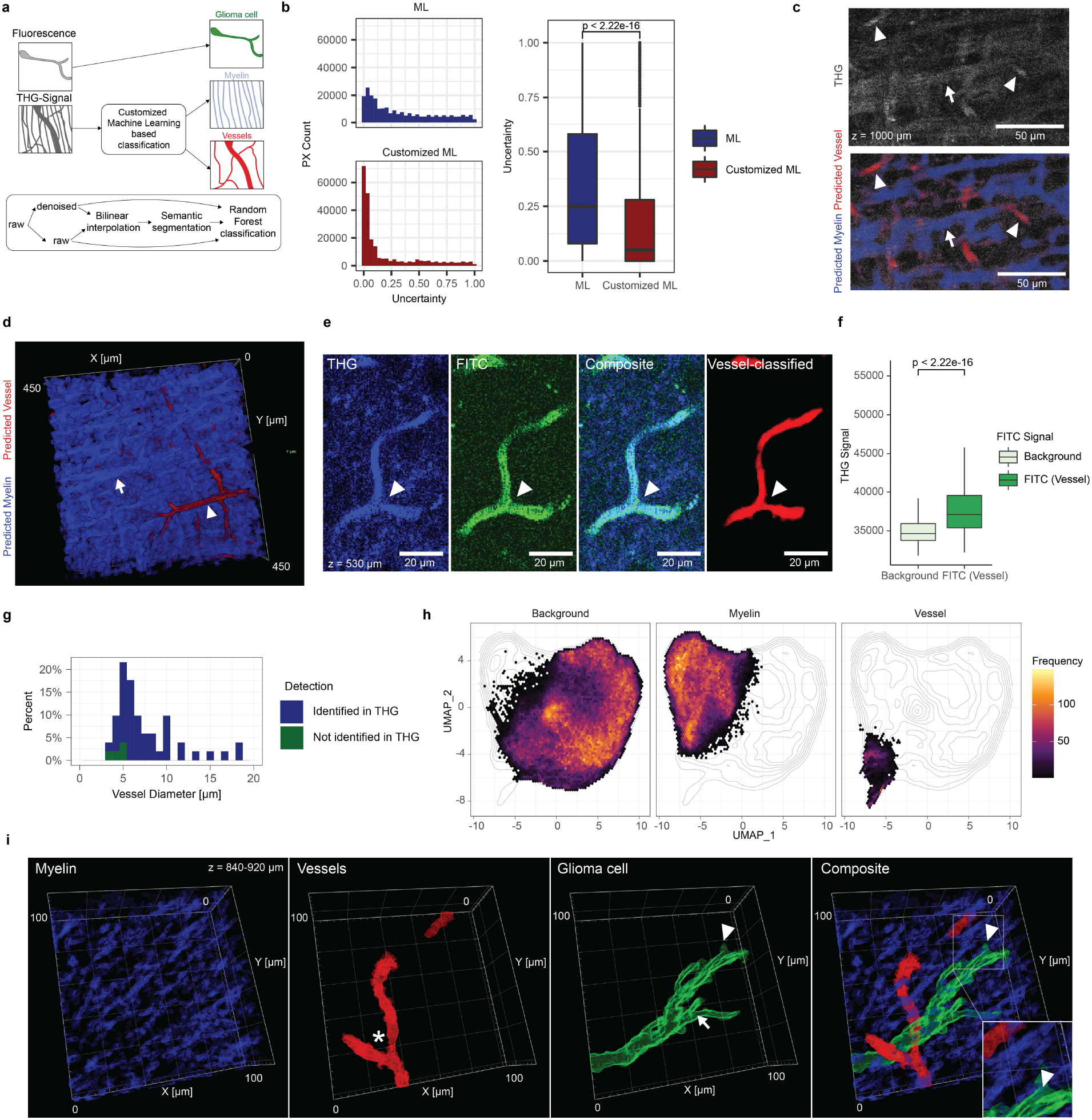
Machine learning-based multicolor imaging of glioblastoma, blood vessels and white matter tracts. a, Scheme for customized machine learning based classification of THG signal into myelin and vessel signal. **b**, Distribution of uncertainty level with machine learning compared with customized machine learning (left) and statistical comparison of uncertainty levels (n = 239568 THG pixels, Mann-Whitney test). **c,** Exemplary image of THG signal without (top) and with (bottom) predicted labels of blood vessels and myelin fibers. Arrowheads indicate blood vessels; arrow indicates myelin fibers. **d,** 3D rendering within the corpus callosum, illustrating the results of the machine learning based classification. Arrowhead indicates vessels, arrow indicates myelin fibers. **e,** Validation of ML-classification for blood vessels with FITC as fluorescent dye colored in green. The arrowheads point at vessels. **f,** Comparison of high and low FITC signal with THG signal (n = 62244 pixels, Mann-Whitney test). **g,** Histogram of measured blood vessels based on their diameter and colored based on their identifcation from THG signal (blue: visible with FITC and in THG signal, green: visible only with FITC, n = 51 vessels). **h,** UMAP based on pixel features that are different between background, myelin and vessels based on the machine learning prediction (n = 100 features). Pixels are colored based on the local frequency of pixels in the dimensionality-reduced space (n = 239568 pixels). **i,** Close-up 3D rendering of a single glioblastoma cell (green) and its surrounding microenvironment (vessel in red, myelin in blue). The asterisk points at a vessel branching points, the arrow at a TM branching point and the arrowhead at a glioblastoma small process. Gamma values were adjusted for 3D-visualization in **d**, **i**.

We further corroborated that the THG signal carries distinct information in its pixel distribution of blood vessels and myelinated white matter tracts using a dimensionality reduction analysis using uniform manifold approximation and projection (UMAP) (Fig. 4h, see Methods). Comparing our customized machine-learning workflow to the standard machine learning pipeline, we could observe that the amount of pixels classified with high uncertainty significantly decreased (Fig. 4b). Lastly, we validated the classifier with human annotation (Extended Data Fig. 4a). Taken together, this workflow enables us to analyze both tumor cells and their microenvironment of blood vessels and myelinated axonal tracts at the same time using a single fluorescence detection channel (Fig. 4f, Extended Data Fig. 4b). The near diffraction-limited resolution of Deep3P allows to clearly identify TMs and the even finer class of neurite-like structures called small processes^3^ in the corpus callosum (Fig. 4i).

## Dynamic deep brain investigation of glioblastoma invasion

Our Deep3P imaging workflow and analysis revealed a number of distinct glioblastoma invasion and colonization patterns within the deep cortex and white matter as well as microenvironmental changes in the white matter *in vivo* that were enabled by the near diffraction-limited resolution and deep imaging depth.

In the corpus callosum, a majority of glioblastoma cells were aligned with myelinated axonal tracts, both with respect to their cell somata and their TMs (Fig. 5a-c) indicating distinct morphological adaptations of glioblastoma cells in the cortex and corpus callosum, depending on the tumor microenvironment. Statistical comparisons of glioblastoma cell and TM directionality analyses revealed a significant correlation of tumor cell directionality with its microenvironment (Fig. 5c). Cell polarity in the cortex showed no clearly structured direction, while in the corpus callosum more than 60 percent of cells grew parallel to the myelin fibers in an angle smaller than 30° (Fig. 5c). Interestingly, a subpopulation of glioblastoma cells was found to be not aligned with the corpus callosum fibers (Fig. 5a-c). Next, we used Deep3P and its stable time-lapse deep brain imaging ability to analyze how glioblastoma cells could invade from the deep cortex into the distinct microenvironment and white matter-rich structure of the corpus callosum (Extended Data Fig. 1a). We found that there is a significant enrichment of perivascular invasion of glioblastoma cells to enter the corpus callosum as compared to perivascular invasion prevalent within the cortex (Fig. 5d-e). An average of 60 percent of glioblastoma cells use vessels as a track to enter the corpus callosum while less than 40 percent of glioblastoma cells in the cortex show a perivascular migration pattern. This illustrates how tumor cells are able to adapt their invasion strategy making use of blood vessels to invade the white matter. Furthermore, Deep3P revealed two vascular mechanisms within the corpus callosum that allowed an invasion that is orthogonal to the myelinated axonal tract direction. First, we discern that glioblastoma cells could extend their TMs to attach to vessels, revealed by the high spatial resolution afforded by the use of our AO (Fig. 5f-h). Using this anchor point, glioblastoma cell somata can then translocate their soma towards the vessel as an invasion mechanism. Furthermore, we could also observe perivascular invasion as an alternative vascular route (Fig. 5f-h). Most vessels to which tumor cells attached to had vessel diameters typical of capillaries^32^ (Fig. 5i). We confirmed the structural relationship between TMs, glioblastoma cell somata and vessels with three-dimensional electron microscopy reconstructions in a patient-derived xenograft model, validating the ability of Deep3P to uncover novel biology at high resolution (Extended Data Fig. 5).

**Fig. 5.**
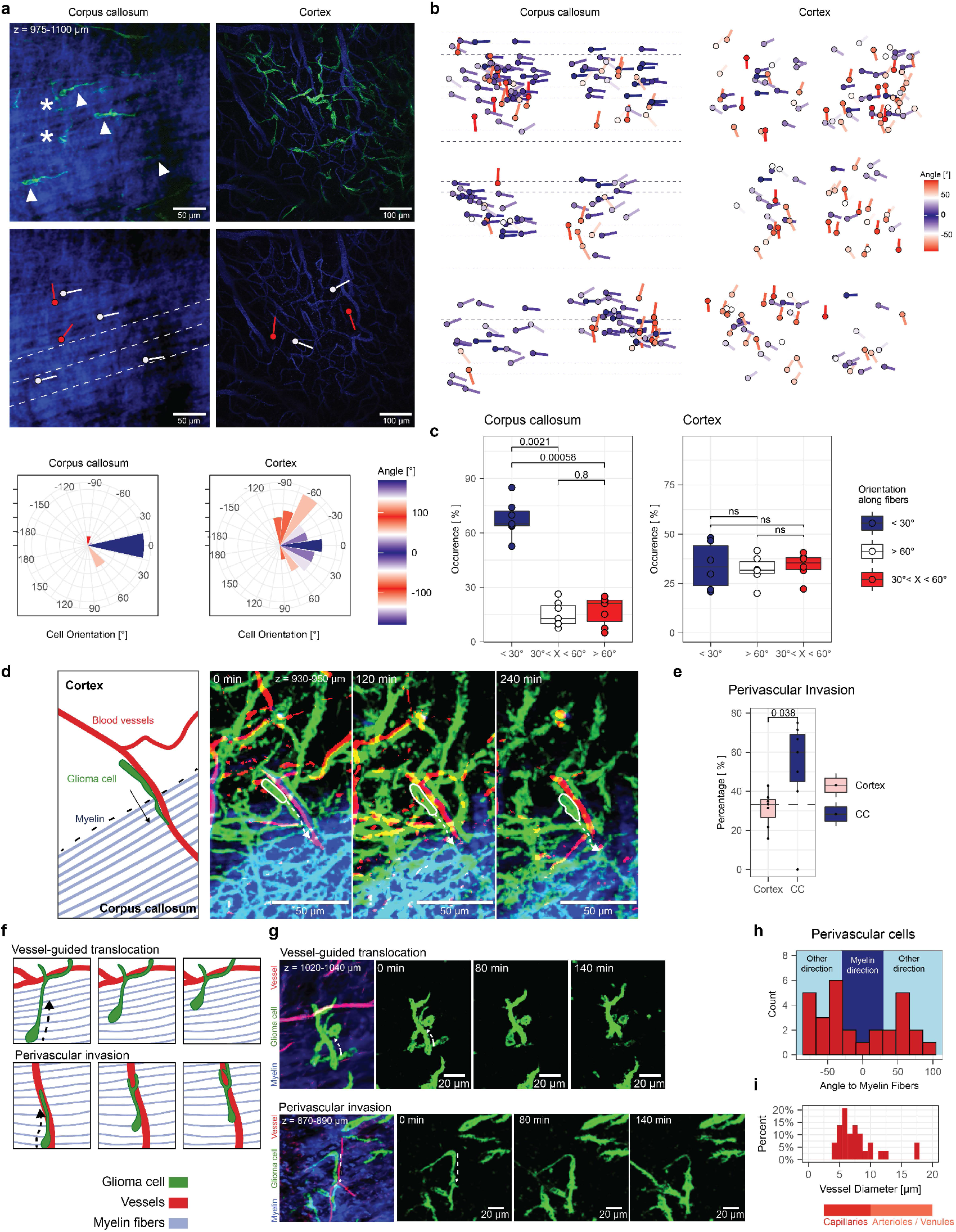
Glioblastoma cell polarity and vascular invasion patterns into and in the corpus callosum. a, Top: Maximum intensity projections of regional overviews of glioblastoma cells in the corpus callosum compared to the cortex are shown. The arrowheads point to exemplary cells that are parallel to the myelin fibers (angle < 30°), the asterisks show exemplary orthogonally oriented glioblastoma cells. Middle left: Corresponding THG signal only. The dashed lines indicate the direction of the myelin fibers, white structures represent exemplary glioblastoma cells aligned with white matter tracts as shown above, the red structure represent exemplary almost orthogonally orientated glioblastoma cells. Middle right: Vessel channel showing the location and direction of four exemplary glioblastoma cells in the cortex represented by the red and white structures. Bottom: Rose plot of tumor cell directionality is shown of the regions above in the corpus callosum and cortex, respectively (n = 67 cells from n = 2 experiments). **b,** Directionality analysis of tumor cell regions in the corpus callosum (left) and in the cortex (right) (n = 355 cells from n = 12 experiments in n = 10 mice in 2 PDX models (S24 and T269)). Dashed lines indicate the direction of the corpus callosum myelinated fibers. **c,** Comparison of predominant angle direction of cell polarity in the corpus callosum and in the cortex (n = 211 glioblastoma cells in the corpus callosum, and n = 164 glioblastoma cells in cortex from n = 13 experiments in n = 11 mice in 2 PDX models (S24 and T269)). Mann-Whitney test). **d,** Maximum intensity projection of an exemplary glioblastoma cell that uses vessels to invade into the corpus callosum. The white circle indicates a glioblastoma cell soma, other glioma cells also visible in the background (green). The dashed line indicates the direction of glioblastoma cell invasion along the blood vessel. Vessels (red), tumor cells (green) and myelinated fiber (blue) are shown as probability maps. **e,** Percentage of glioblastoma cells showing perivascular invasion into the corpus callosum as compared to perivascular invasion within the cortex (n = 206 glioblastoma cells from n = 14 experiments, Mann-Whitney test). **f,** Schematic drawing of how tumor cells use orthogonal vessel migration or perivascular invasion. **g,** Example of how tumor cells use orthogonal vessel migration or perivascular invasion. The dashed line indicates the direction of glioblastoma cell invasion. Maximum intensity projections of vessels (red) and tumor cells (green) are shown as probability maps and the signal of the corpus callosum (blue) underwent denoising post-processing. **h,** Histogram of angles between perivascular cells and myelin fiber orientation. (n = 29 glioblastoma cells from n = 7 experiments in 5 mice in 2 PDX models (S24 and T269)). **i,** Histogram of blood vessel diameter of perivascular cells (n = 29 glioblastoma cells from n = 7 experiments in 5 mice in 2 PDX models (S24 and T269)).

Interestingly, these invasion patterns resemble migration patterns of oligodendrocytic and astrocytic precursor cells during neurodevelopment^26, 27^. This shows that not only cell-intrinsic mechanisms of neural precursor cells are hijacked by glioblastoma cells for invasion but also that their microenvironmental dependencies are phenocopied.

Subcellular dynamic behavior of TMs including branching, protrusion, and retraction, previously only described in the superficial cortex^3^, were observed in both deep gray and white matter (Fig. 6a-b, Extended Data Fig. 6). Interestingly, branching of TMs was reduced within the corpus callosum, potentially indicative of a more directed movement pattern of TMs along white matter tracts as compared to a scanning behavior of TMs prevalent in the superficial layers of the cortex^3^. Furthermore, three distinct invasion patterns of locomotion, branching migration and translocation could be detected within the corpus callosum, thanks to the superior spatial resolution and depth penetration of Deep3P (Fig. 6c-d, Supplementary Video 4). These migration mechanisms reflect conserved neuronal mechanisms of invasion that can be seen during neuronal development^3, 33^. Interestingly, this is congruent with previous observations in the superficial layers of the cortex^3^. However, in contrast to invasion within the cortex branching migration within the white matter is significantly reduced in line with reduced branching behavior of the TMs (Fig. 6d). It will be important to characterize the distinct molecular mechanisms of these three invasion phenotypes in their microenvironmental niches of the gray and white matter which is now possible via Deep3P. While these mechanisms are phenotypically distinct, their invasion speed did not significantly differ (Fig. 6e).

**Fig. 6.**
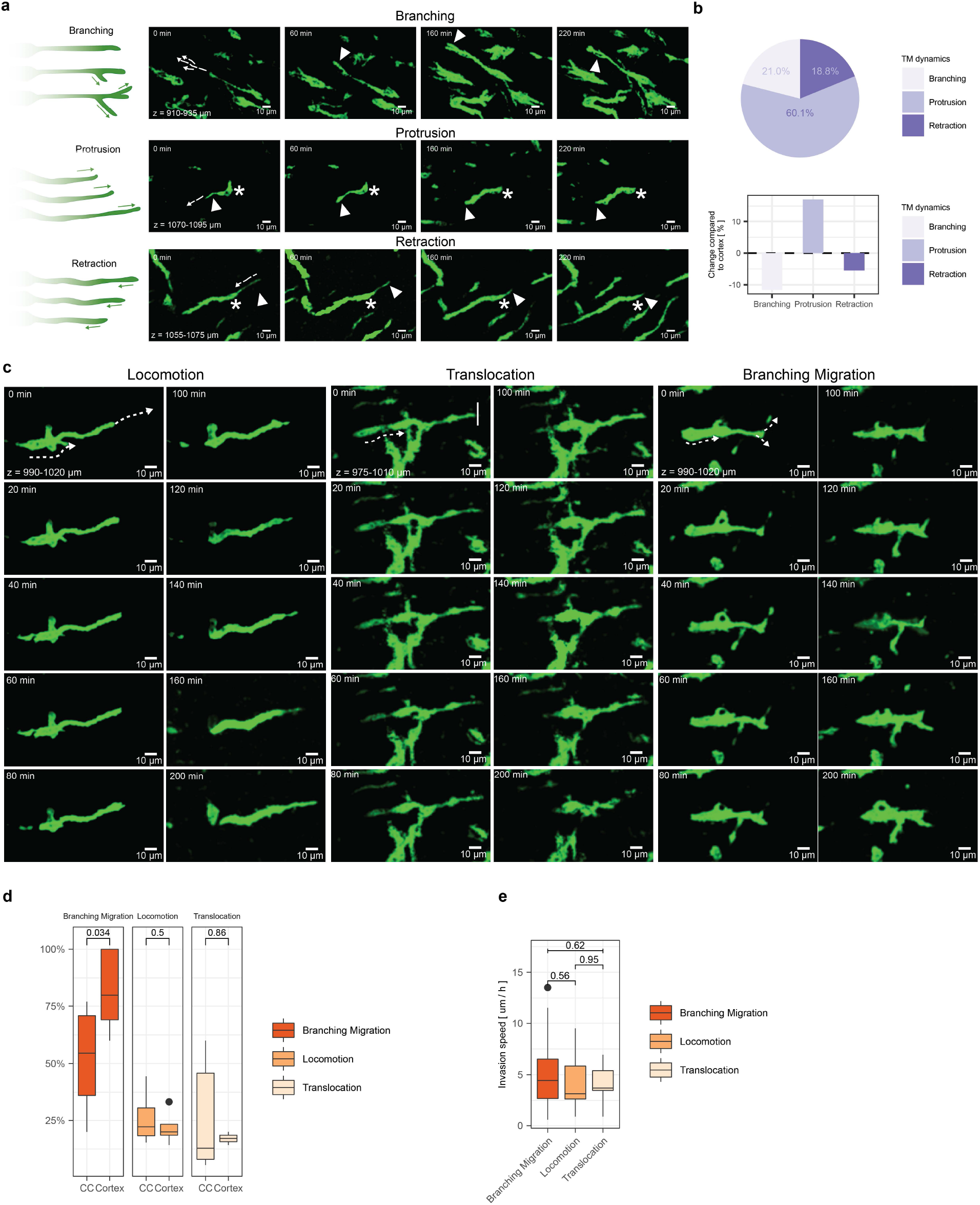
TM dynamics and neural invasion mechanisms in the corpus callosum. a, Representative maximum intensity projection time-lapse images showing TMs in the corpus callosum that use branching, protrusion, or retraction. Asterisks point at the glioblastoma cell somata, arrowheads point towards the tips of the TMs of interest. Dashed arrows show the direction of the TM dynamic. **b,** Distribution of TM dynamics in the corpus callosum and comparison with cortex using branching, protrusion or retraction (n = 163 cells from n = 14 datasets in 12 mice in 2 PDX models (S24 and T269)). **c,** Representative maximum intensity projection time-lapse images of invasion phenotypes in corpus callosum showing locomotion, translocation and branching migration. The dashed arrows show the invasion direction. **d,** Distribution of invasion phenotypes in the corpus callosum compared to the cortex (n = 117 cells from n = 7 experiments in 6 mice, Mann-Whitney test). **e,** Speed comparison of invasion phenotypes in corpus callosum (n = 45 cells from n = 7 experiments in 6 mice, Mann-Whitney test). Images were post-processed as probability maps and using the ‘‘smooth” function in ImageJ/Fiji in **a**, **c**.

### Glioblastoma network formation and tumor cell proliferation in the white matter

In addition to gap junction-coupled tumor-tumor networks that were so far exclusively described in the gray matter^3, 6, 15, 34–36^, we could also observe how tumor-tumor networks in the white matter are formed (Fig. 7a-c, Extended Data Fig. 7a, Supplementary Video 5). In contrast to tumor-tumor networks in the gray matter, these tumor-tumor networks are to a majority aligned with the white matter fiber direction of the corpus callosum as compared to the cortex (Fig.7b-c). This illustrates how tumor network formation respects the anatomical boundaries and integrates into the peculiar microenvironment of the corpus callosum (Fig. 7a-c). In addition to investigating invasion and tumor network formation, Deep3P could also be used to characterize the specialized patterns of glioblastoma cell division qualitatively and quantitatively within the deep, native microenvironment of the corpus callosum (Fig. 7d, Extended Data Fig 7b, Supplementary Video 6).

**Fig. 7.**
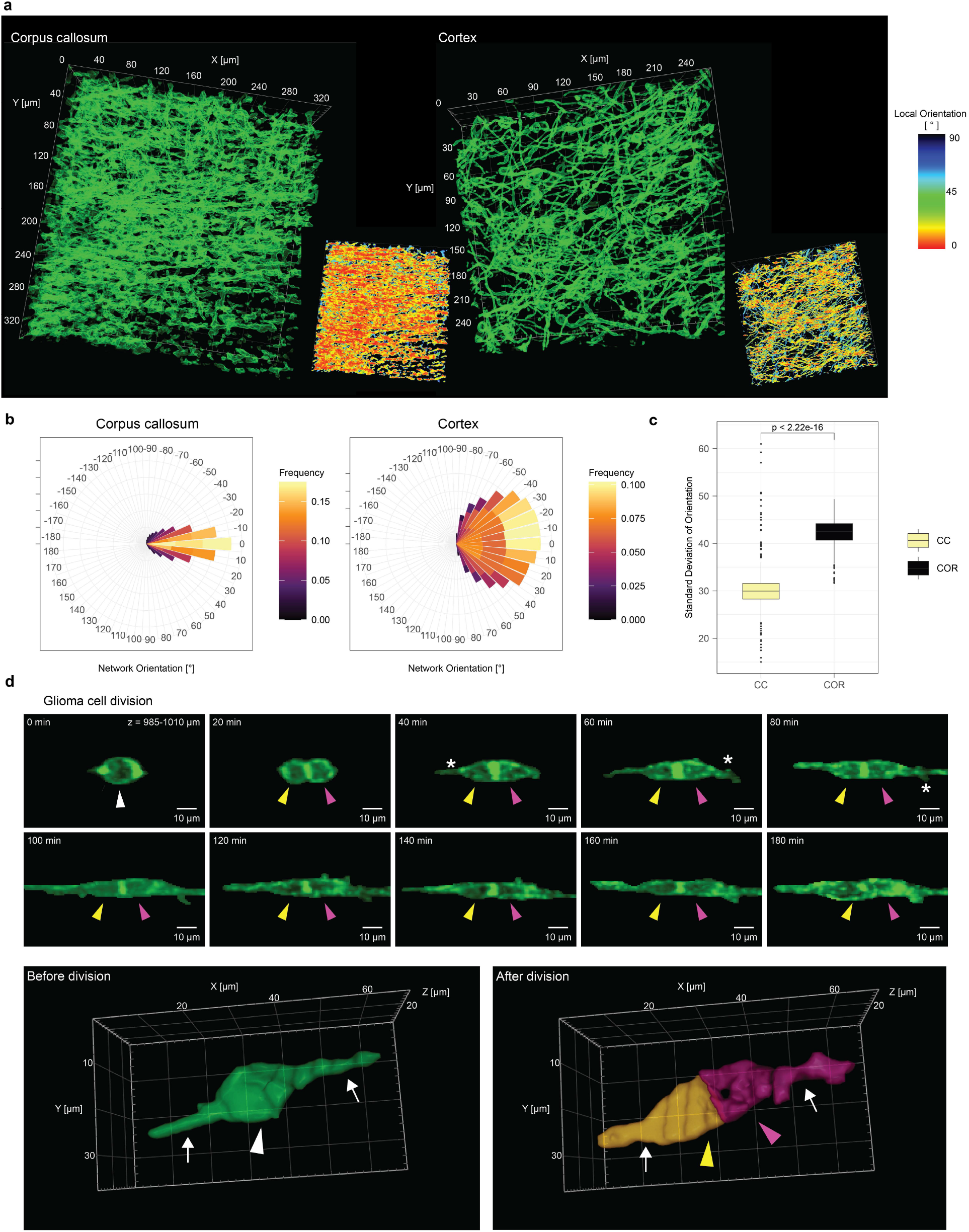
Tumor-tumor network formation and glioma cell proliferation in cortex and corpus callosum. a, Brain tumor networks in the corpus callosum (left) and in the cortex (right) shown as 3 D renderings in green. Corpus callosum and cortex imaging were performed with 3PM and 2PM, respectively. Each network is visualized as network orientation in the bottom right corner. The network is colored based on the local orientation. Corpus callosum imaging depth: z = 840-950 µm. **b,** Rose plot of overall orientation of the tumor cell network in the corpus callosum (n = 271985 local network orientation values from n = 6 brain tumor networks from n = 5 mice). **c,** The standard deviation of tumor network orientation is compared in the corpus callosum to the cortex. (n = 692 slices from n = 6 brain tumor networks from n = 5 mice, Mann-Whitney test). **d,** Top: Maximum intensity projection time-lapse imaging of glioblastoma cell division in the corpus callosum at an imaging depth of 985-1010 µm. White arrowhead: Glioblastoma cell before cell division. Yellow and purple arrowhead: Daughter glioblastoma cells after cell division. The asterisk points at a newly grown TM after cell division. Post-processed with denoising and “clear outside” function in ImageJ/Fiji. Bottom: 3D rendering of another cell in the corpus callosum before and after cell division. The arrowheads point to the somata of the cell before division and the two daughter cells after division, the arrows point to the TMs (n = 18 cell divisions from n = 7 experiments in 6 mice in 2 PDX models (S24 and T269)). Gamma values were adjusted for 3D-visualization in **a**, **d**.

### White matter disruptions during early glioblastoma colonization

Lastly, we investigated the role of glioblastoma invasion and early colonization in the corpus callosum on myelinated fiber tracts. Evaluating white matter integrity during glioma infiltration with clinical imaging such as magnet resonance imaging (MRI) would be a useful bioimaging marker for estimating whole-brain colonization of glioma. It has been proposed that diffusion tensor imaging (DTI) enables imaging of early tumor infiltration into the corpus callosum^37^. However, a comparison to the ground truth of tumor invasion and an exact characterization of tumor density was not possible. To investigate this phenomenon during early glioblastoma invasion and colonization, we longitudinally performed MRI scans of tumor infiltrated mouse brains (Fig. 8a) and analyzed the fractional anisotropy of different regions in the corpus callosum. Over weeks, before and after tumor infiltration, no changes in fractional anisotropy values were observed (Fig. 8b). To investigate whether this observation can be explained on a microscopic scale, we used Deep3P to investigate the effect of tumor infiltration on THG signal in tumor infiltrated regions of the corpus callosum (Extended Data Fig. 8a).

**Fig. 8.**
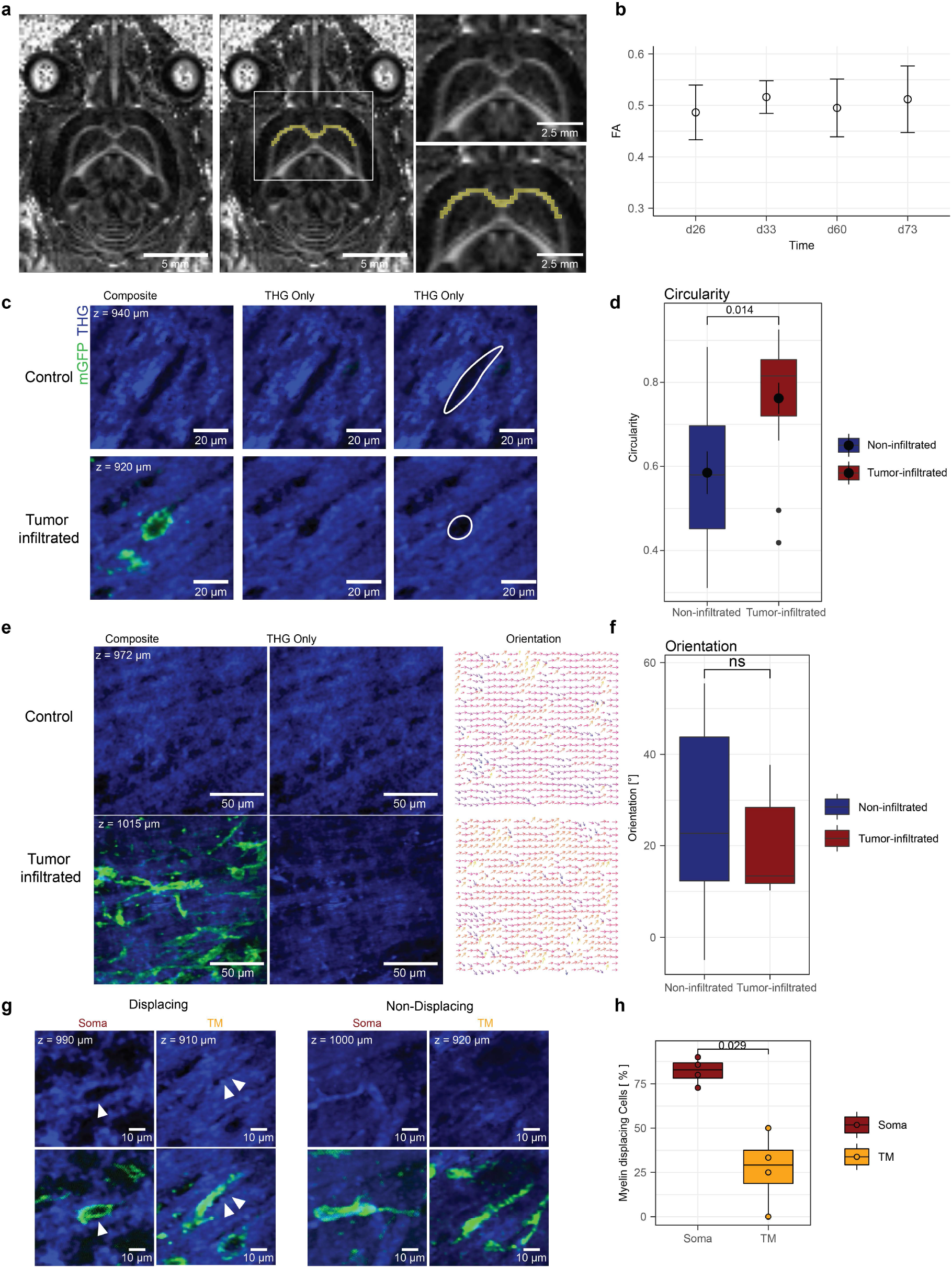
White matter reactivity in the corpus callosum upon glioblastoma invasion and colonization. a, MRI fractional anisotropy map of a horizontal section in a glioma-infiltrated mouse brain (left), with labels on corpus callosum (yellow). Right: Zoom in on corpus callosum, shown is the body part of the CC. (PDX model: S24) **b,** The mean FA-value of mice at multiple time points in the body part of the corpus callosum (d26: 26 days after injection (n=4), d33 (n=8), d60 (n=10), d73 (n=9)).**c,** Regions with and without tumor infiltration were analyzed in the corpus callosum. mGFP and source THG signal were post-processed with denoising. **d,** Morphological parameter (circularity) of tumor-infiltrated holes and non-infiltrated holes (n = 27 cells from n = 3 experiments in 3 mice in 2 PDX models (S24 and T269), Mann-Whitney test). **e,** Left: Corpus callosum fibers in tumor-infiltrated regions compared to non-infiltrated regions. Images were post-processed with denoising. Right: Orientation plot of fibers is shown. **f,** Comparison of orientation of brain tumor free tissue with tumor-infiltrated regions. (n = 29 sample regions from n = 8 mice (S24 and T269), Mann-Whitney test). **g,** White matter displacing (left) and non-displacing (right) glioblastoma cell somata and TMs. Arrowheads show tumor-infiltrated-holes. Images were post-processed with denoising. **h,** Comparison of portion of displacing TMs to displacing glioblastoma cell somata (n = 33 glioblastoma cell somata and n = 22 TMs from n = 4 mice in 2 PDX models (S24 and T269), Mann-Whitney test).

We were able to quantify changes in the white matter tracts that were disrupted by glioblastoma cell invasion in the early stages of glioblastoma colonization using our customized machine learning approach of white matter classification (Fig. 8). We found that the shape of THG discontinuities could potentially serve as an imaging biomarker of early glioblastoma cell invasion into white matter tracts with potential clinical-translational implications. While glioblastoma cells in the white matter significantly showed an increased roundness of their cells, the resident cells of the tumor microenvironment were more oval-shaped in the corpus callosum (Fig. 8c-d). However, the white matter tract directionality did not change significantly in the early stages of glioblastoma colonization (Fig. 8e-f), potentially also explaining difficulties of modern clinical imaging to delineate white matter glioma infiltration^38^. Additionally, we observed that TMs, the fine neurite-like protrusions, displaced white matter tracts significantly less than their glioblastoma cell soma (Fig. 8g-h). This might indicate that the TMs are used to scan and invade the microenvironment without causing major disruptions of its myelinated microenvironment. Taken together, these observations could serve as ground truth for further development of microscopy approaches and high-resolution clinical imaging to detect early glioblastoma invasion.

## Discussion

In summary, we developed a bespoke intravital imaging workflow that uses 2PM and 3PM together with AO enabling deep brain tumor imaging up to a depth of 1.2 mm in the living mouse brain. This methodology opens up new avenues to study the cancer neuroscience of brain tumors beyond superficial cortex layers in the pathophysiologically highly relevant white matter.

Compared to previous proof-of-principle technical work in 3PM and AO^18, 20, 22, 39, 40^, we optimized and advanced these technologies for longitudinal intravital measurements at the sub-cellular scale in >1 mm depth and over several hours for samples with low SNR. Instrumental to this was the diligent optimization of imaging parameters, conditions and overall workflow (see Supplementary Note 1), as well as the addition of an artificial intelligence-driven image restoration workflow adapted to the distinct noise sources of 3PM to substantially increase image SNR and thereby keep overall excitation light photoburden minimal. Overall, this workflow significantly increased the robustness of 3PM ensuring to study white matter glioblastoma biology non-invasively *in situ*. Lastly, we used a bespoke machine-learning algorithm to separate the THG signal and thereby to clearly distinguish most blood vessels and white matter tracts, enabling simultaneous volumetric time-lapse imaging of tumor cells and their microenvironment. At the moment, the main disadvantage of our Deep3P methodology is the relatively slow acquisition speed, due to constraints associated with laser power, pulse energy and sample physiology^20^, as well as AO optimization time overheads, which limits the overall volumetric FOV that can be monitored over time. Yet, the introduction of efficient 2P pre-screening as well as the comparably fast modal-based AO procedure^22^ ensure that biologically interesting brain regions can be effectively and non-invasively imaged over time.

The effectiveness of our Deep3P approach is demonstrated by its ability to track highly dynamic subcellular and cellular processes of glioblastoma in white and gray matter. This approach uncovered different adaptations of glioblastoma cell morphology, directionality and invasion phenotypes in the corpus callosum as compared to the cortex. For these findings, the superior depth penetration, spatial resolution as well as image SNR was of paramount importance. Our approach also allows for the intravital investigation of tumor biology including proliferation, invasion, and therapeutic resistance across different brain regions and brain tumor entities. Further, the machine learning-based separation of the THG signal allowed the discrimination of blood vessels and the myelinated axons from the raw image data. This analysis approach can be in principle also generalized and adapted to other non-linear microscopy techniques that capture label-free image contrasts of different biological structures.

The understanding of invasion mechanisms deep in the brain and their relationship to the microenvironmental niches of blood vessels and myelinated fibers are fundamental to glioblastoma biology and potential therapies. We find not only a novel mechanism of vascular translocation that is mechanistically distinct to the perivascular invasion route. In addition, we observe an enrichment of migration along the vascular route into the corpus callosum with different invasion mechanisms prevalent within the corpus callosum. While the route of perivascular migration has been previously described^16, 17^ and association with a breakdown of the blood-brain barrier neither vascular translocation at all nor perivascular migration in the corpus callosum was described yet. Interestingly, we observe these phenomena in the very early stages of glioblastoma colonization where the blood-brain barrier is still intact. Taken together, these are fundamental in our understanding of glioblastoma biology.

In addition to its utility in brain tumor imaging, we envision that our newly developed integrated microscopy and analysis workflow allows for the integration of fluorescent cell state indicators and other subcellular fluorescent reporters with low SNR to simultaneously characterize biological behavior as well as cellular and molecular cell states in the future. Furthermore, this method enables the development of novel correlative technologies deep in the brain, as previously demonstrated for correlative light and electron microscopy, allowing for in-depth characterization of intravital cellular behavior and ultrastructure in a non-invasive manner.

Apart from patient-derived brain tumor models, we believe that our 3PM and analysis workflow can be straightforwardly applied to other disease models such as other brain tumors, extracranial tumors, demyelinating diseases, including imaging of the spinal cord and, as well as across various model organisms to investigate (sub)cellular structure and (patho)physiology with minimal invasiveness and samples with low SNR. Further red-shifted 3PM and/or its combination with wavefront-shaping approaches could lead to even increased imaging depth beyond the corpus callosum to truly enable whole-brain imaging of glioblastoma colonization. Integrating feedback microscopy, also utilizing AI to identify regions of interest, might provide a powerful solution for faster and more automated imaging in the future.

Interestingly, our investigation of white matter disruptions revealed only minor changes of the overall structure in myelinated white matter tracts. These near diffraction-limited microscopical findings further explain difficulties of clinical imaging^38^ to diagnose the earliest stages of glioblastoma colonization in clinical settings.

Overall, our approach allows for the investigation of spatiotemporally dynamic brain tumor biology *in vivo* across the gray and white matter, with the potential to further uncover novel cellular and subcellular mechanisms in cancer and its microenvironment. This has important implications for our understanding of brain tumors, opportunities to study these devastating diseases, and the development of new clinical therapies.

## Author contributions

Supervision, R.P., V.V.; Conceptualization, R.P., V.V.; Methodology, S.S., M.S., A.T., E.M., A.K., R.P., V.V.; Software Analysis, M.S., Software Denoising: E.M.; Project Administration, R.P., V.V.; Investigation, S.S., M.S., A.T., R.D., M.D., N.W.; Formal Analysis, S.S., M.S., A.T., M.D., R.D., E.R., S.T., N.W., C.B., J.S., F.T.K.; Resources, S. H., A.G., F. T. K., W.W., T.K., F.W., C.H., R.P., V.V.; Visualization, S.S., M.S., E.R., C.B.; Writing-Original Draft, R.P., V.V.; Writing-Review & Editing, S.S., M.S., A.T., E.M., R.D., M.D., E.R., S.T., N.W., A.G., C.B., W.W., F.W., A.K., T.K., C.H., R.P., V.V..; Funding Acquisition, W.W., F.W., T.K., R.P., V.V.

## Acknowledgements

V.V. received financial support from the German Research Foundation (DFG: VE1373/2-1), the Else Kröner-Fresenius-Stiftung (2020-EKEA.135), the University of Heidelberg (Physician-Scientist-Program, Krebs-und Scharlachstiftung) and the Health + Life Science Alliance Heidelberg Mannheim. M.S., E.R. and S.T. were supported by the Deutsche Krebshilfe/German Cancer Aid (Mildred-Scheel-Scholarship for MD students). W.W., F.W. and V.V. were supported by a grant from the German Research Foundation (DFG: SFB 1389). R.P. acknowledges support from the European Commission (grant no. 951991, Brainiaqs), and the Chan Zuckerberg Initiative (Deep Tissue Imaging grant no. 2020-225346). R.P., A.T. and E.M. were supported by the EMBL. The Northwestern Nervous System Tumor Bank is supported by the P50CA221747 SPORE for Translational Approaches to Brain Cancer. F.T.K. was supported by a grant from the German Research Foundation (# 507778602).

We thank M. Kaiser, Y. Dörflinger, S. Hoppe, M. Schmitt, A. Oudin and V. Baus for technical assistance. We thank F. Blum, T. Rubner, M. Eich, K. Hexel and S. Schmitt from the Flow Cytometry Core Facility of the German Cancer Research center for their assistance in performing FACS experiments. We thank K. Becker for providing support and assistance in animal care and design of animal experiments. We thank the EMBL animal facilities for help and support. We thank the EM Core Facility of the Heidelberg University and the EM and Microscopy Core Facility of the German Cancer Research center for their support. The authors thank the donors and their families who donated through the Nervous System Tumor Bank.

## Materials and Methods

### Experimental models and subject details

All animal studies were performed with adult male NMRI nude mice in accordance with the European Directive on animal experimentation (2010/63/EU) and institutional laboratory animal research guidelines after approval of the Regierungspräsidium Karlsruhe, Germany, the EMBL IACUC and the Animal Welfare Structure of the Luxembourg Institute of Health (protocol LRNO-2017-01). The 3R principles for reducing the number of animals were strictly followed and efforts were made to minimize animal suffering. Animals were scored daily and experiments were terminated in case of weight loss exceeding 10-20%, neurological deficits and signs of pain, tumor size was no termination criterion. For all human tissues, patients have given informed consent and local regulatory authorities have approved (Ethic Committees at the Mannheim and Heidelberg Medical Faculty of the University Heidelberg, protocols (S-206/2005, S-207/2005, S-306/2019, 2018-614N-MA, 2018-843R-MA), the National Committee for Ethics in Research (CNER) Luxembourg (201201/06) and the regionale komiteer for medisinsk og helsefaglig forskningsetikk at the Helse Bergen (protocol 2013/720/REK vest). Informed consent was obtained for all patients who donated their brain post-mortem to the Nervous System Tumor Bank. The protocol was reviewed and approved by the Northwestern University Institutional Review Board (IRB) under study STU00095863. Molecular testing with antibodies against the IDH mutation, ATRX stainings and the Illumina 850k methylation array (Department of Neuropathology, University of Heidelberg) for confirmation of diagnosis were performed.

### Patient-derived primary glioblastoma cell lines and Illumina 850k methylation array characterization

Cultivation of patient-derived tumor cell lines from resected glioblastoma was performed as previously described^6, 15, 34, 41^ in DMEM/F-12 under serum-free, non-adherent, ‘stem-like’ conditions with B27 supplement (12587-010, Gibco), insulin, heparin, epidermal growth factor, and fibroblast growth factor as described previously^15^. The molecular classification of glioblastoma xenograft models used in this study can be found in Supplementary Table 2. To obtain the DNA methylation status at >850,000 CpG sites in all GBC lines, the Illumina Infinium Methylation EPIC kit was used according to the manufacturer’s instructions at the Genomics and Proteomics Core Facility of the German Cancer Research Center in Heidelberg, Germany, as described previously^42^. Glioblastoma cell lines kept under stem-like conditions were transduced with lentiviral vectors for membrane-bound GFP with the pLego-T2-mGFP construct based on Dondzillo et al.^43^.

Regular FACS sorting of transduced cells was performed with FACSAria Fusion 2 Bernhard Shoor or FACSAria Fusion Richard Sweet and the BL530/30 filter was used for FACS-sorting GFP.

### Surgical procedures

Surgical procedures were performed as previously described^6, 15^. Cranial window implantation in mice was done similarly to what we had previously described with small modifications, including a custom-made titanium ring for painless head fixation during imaging and an asymmetric placement of the window above the sinus, allowing optimal imaging accessibility of the corpus callosum on one hemisphere. 50.000-100.000 tumor cells were stereotactically injected at a depth of 500 μm into the mouse cortex.

### Histological analysis of organoid-based patient-derived orthotopic xenograft models

Patient-derived orthotopic xenografts were derived by intracranial implantation of glioma organoids as described in Oudin et al.,2021^44^. PDOXs were assessed at the histopathological and molecular levels as described in Golebiewska et al, 2020^45^. Invasion of human glioma cells through corpus callosum was assessed by immunohistochemistry with antibodies against human-specific nestin (abcam, ab6320, 1:500) and/or vimentin (Themo Fisher Scientific, Mab3400, 1:200) on coronal 4-8µm sections from paraffin-embedded brains. Primary antibodies were incubated overnight at 4°C or 3 hours at room temperature, followed by 30 min incubation with secondary antibodies. Signal was developed with the Envision+ System/HRP Kit in 5–20 min (K4007, Agilent/Dako).

### Spatial transcriptomics analysis of cortex and corpus callosum

Spatial sequencing data as well as imaging data was downloaded from http://molecularatlas.org based on Ortiz et al., 2020^46^. Data was processed using Seurat^47^ and clusters were defined using its FindClusters function. Based on the associated histology images, clusters were assigned to either corpus callosum / white matter, cortex / gray matter, or other brain regions. Clusters were then visualized based on the assignment and using a Uniform Manifold Approximation and Projection (UMAP) in Seurat.

### Macroscopic and microscopic histological analysis of glioblastoma patients

The postmortem cohort was collected as described in Drumm, et al.^1^. Briefly, at the time of brain sectioning, portions of key brain and spinal cord regions, as well as extra sampling of tumor, were collected for histologic processing as paraffin-embedded tissue blocks. Tissue sections of corpus callosum were routinely collected whether gross evidence of tumor involvement was present or not. Each section of corpus callosum was then stained with hematoxylin and eosin and examined for the presence of migrating glioma cells.

Standard immunohistochemistry analysis was performed on patient autopsy specimens to visualize IDH1R132H (Dianova DIA H09, 1:150) and nestin (Abcam ab22035, 1:750). To achieve this, four-micrometer thick sections of FFPE tissue on charged slides were baked in the oven at 60C before being deparrafinized and re-hydrated. Antigen retrieval was performed using a pH6 retrieval buffer (Biocare Reveal). Slides were cooled to room temperature and washed in TBS before neutralizing endogenous peroxidase (Biocare Peroxidase 1). Slides were then treated with a serum-free casein background block (Biocare Background Sniper) before incubation in a 10% goat serum block for 60 minutes at room temperature. Primary antibody was then added to the slides for overnight incubation at 4C. After incubation, slides were washed well with TBS-T before incubating in HRP polymer (Biocare MACH 4 Universal HRP Polymer). Finally, reaction products were visualized with DAB (Biocare Betazoid DAB Chromogen Kit). Slides were then counterstained with hematoxylin, dehydrated and mounted with xylene-based mounting media.

### Two-photon microscopy

For 2PM imaging, chronic cranial window surgeries were performed on adult male NMRI nude mice (Charles River and Janvier) and patient-derived tumor cells were injected into the cortex as described previously. 2PM was first performed three weeks after surgery using a TriM Scope II microscope (LaVision BioTec GmbH) equipped with a pulsed Ti:Sapphire laser (Chameleon II ultra; Coherent). A 16×, 0.8 NA, apochromatic, 3 mm working distance, water immersion objective was used. Before imaging, TRITC dextran (fluorescent conjugated tetramethylrhodamine isothiocyanate-dextran, 500.000 g/mol) was diluted in 0.9% NaCl-solution at 10 mg/ml and 100 µl injected intravenously into the tail vein for vessel signal. GFP and TRITC were imaged using 960 nm wavelengths. Low-noise high-sensitivity photomultiplier tubes were used for fluorescence emission detection. For the anesthesia, isoflurane gas was diluted in 100% O2 to a concentration between 5% (for anesthesia induction) and 0.5-1.5 % (for maintenance of anesthesia). The breathing rate of the mice was monitored and body temperature was kept stable at 37° using a heating pad. Eye cream was applied before anesthesia. The tumor growth was regularly screened using 2PM. To allow easier identification of tumor regions suitable for 3PM corpus callosum imaging, whole brain tile scans were acquired and mapping of the vessel structure was performed in order to allow faster detections of the regions during 3PM.

For whole brain tile scan imaging, repetitive stacks of 2PM images were acquired with a field of view of 694 µm x 694 µm. Images were taken down to a depth of approximately 600-700 µm, with a z-step size of 10-20 µm. Subsequently, stacks were stitched together resulting in an overall field of view of up to 4.3 x 4.3 mm x 0.7 mm using the Fiji^48^-based stitching plugin^49^.

### Three-Photon Microscopy

The core hardware of the multiphoton 3P microscope with adaptive optics has been described in detail in previous work^22^. In the following, a brief summary of the instrumentation is provided emphasizing any differences from the details given in Streich et al.^22^.

Over the course of the study, two excitation sources were used in this work. One of them is a Spectra-Physics TOPAS tunable non-collinear optical parametric amplifier generated ∼60 fs pulses centered at 1300 nm with a repetition rate of 400 kHz, pumped by a 16W Spectra-Physics Spirit. The near-IR pulses from the TOPAS were pre-compensated for dispersion by a homebuilt single prism (N-SF11) pulse compressor^50^. The maximum power at this wavelength and repetition rate was ∼400 mW, resulting in a maximum available power under the objective of 35mW. A Pockels cell was used for rapid modulation of the laser power during image acquisition. Furthermore, we also employed a Class 5 Photonics White Dwarf WD-1300-dual laser. The 1300 nm channel used here provided a maximum of over 5 Watts at a repetition rate of 1 MHz. In addition to the Pockels cell, a reflective optical density filter with a static OD=0.8 attenuation was used to adapt the power range of the White Dwarf, yielding a maximum of 100 mW after the objective. Dispersion pre-compensation was done by an internal module in the White Dwarf which yielded 100 fs pulses after the objective. For fluorescence detection in the green channel, we switched to an uncooled H10770PA-40. The adaptive optics module used an ALPAO DM97-15 continuous membrane deformable mirror, relying on the factory calibration of the mirror for the Zernike-to-Control Matrix.

Images acquired with “full AO” optimization are optimized using the same metric and iterative procedure discussed in Streich et al.^22^ where the fluorescence intensity is measured and optimized directly on mGFP labeled cell in the mouse brain when not specified otherwise. The metric was found to also produce good enhancement when optimized on THG signal, enabling optimization in regions where no labeled cells are available. The typical optimization procedure involved two iterations of Zernike modes 3 through 21, with five amplitudes explored per iteration. With frame rates typically > 1Hz, the entire process is completed in less than 3 minutes even in noisy regions. In both AO optimization as well as imaging, a maximum pulse energy at the focus of 2nJ was never exceeded^51^. Even after a four-hour time-lapse imaging session, only mild bleaching was observed (volume acquisition times range from 6 to 12 minutes which corresponds to a 30% to 60% duty cycle for light exposure since a volume is acquired every 20 minutes, though the time spent on each individual plane is at most 5% of that). The correction collar of the objective was set to 1 mm imaging depth for most acquisitions.

### *In vivo* deep brain imaging

For the initial induction of the anesthesia, nude NMRI mice were placed into a closed box and exposed to 6% isoflurane diluted in oxygen. For the maintenance of the anesthesia, a concentration of 1.6-2% isoflurane in oxygen was used, depending on the mouse’s breathing rate. The targeted breathing rate was between 70 and 90 bpm. Once anesthetized, eye cream was applied and mice were positioned on a small animal physiological monitoring system (ST2 75-1500, Harvard Apparatus), which allowed the maintenance of a stable body temperature at 37.5 °C and monitoring of the breathing rate.

For *in vivo* imaging experiments, animals were head-fixed using a customized headbar and complement holder and the cranial window was aligned so that the objective was placed above the part of the window that is centered slightly next to the sinus (where corpus callosum is anatomically the highest). Acquisition parameters for deep brain imaging are summarized in Supplementary Table 3. Here, we note that imaging depth was reported as raw axial translation of the objective. Taking the refractive index difference between the coverslip, immersion media and various brain tissues into account^18^, the actual imaging depth is likely around 5–10% larger than the values reported. For validation of the THG vessel signal, FITC dextran (fluorescein isothiocyanate-dextran, 2M g/mol) was diluted in 0.9% NaCl-solution at 10 mg/ml and 100 µl injected intravenously into the tail vein.

### Resolution estimation of 3PM

For estimation of image resolution used in comparing images with and without adaptive optics correction, the decorrelation analysis Fiji plugin was used^52^. The calculation was performed for all slices of the green channel in 5 different volume stacks. Slices that were clear outliers (order of magnitude off) or where the calculation failed (decorrelation curves had no maxima) were excluded and a mean and standard deviation for the stack resolution was calculated. The corresponding resolution ratio per volume was calculated with a standard deviation using error propagation. A weighted average of five such volumes was calculated with the corresponding standard deviation.

### 3PM deep learning-based denoising

#### Network architecture

In our approach, we designed a 3D version of the classic U-Net architecture^53, 54^ (see Ext. Data Fig. 2a).The architecture is composed of an encoder-decoder structure connected through skip connections. The encoder network contains 5 hierarchically organized encoder blocks. Each block consists of two consecutive units, including a 3D convolutional layer, a leaky ReLU activation layer, and a group normalization layer. In parallel, the decoder network is made up of 5 blocks with similar layer configurations and a nearest-neighbor upsampling layer at each level. The number of feature maps in the first level of the encoder network is set to 16, and then increases geometrically at each level. The feature maps from the encoder blocks are passed to the corresponding decoder blocks through the skip connections, allowing the combination of high-level features from the encoder with the semantic features from the decoder. This enables the U-Net architecture to effectively incorporate both local and global information in the reconstruction, making it well-suited for image restoration tasks.

#### Self-supervised denoising

In this study, we employed the Noise2Void strategy for self-supervised denoising of 3PM data, which allows for the training on the same data to be denoised^24^. This approach relies on the assumption that the noise is independently generated for each pixel, which is valid for the dominant noise sources in 3PM such as Poisson shot noise and Gaussian readout noise. During the training, a random subset of pixels, referred to as “blind spots”, are masked in the 3D input data and the network is optimized to predict the values of these pixels. This masking forces the prediction to be based solely on the surrounding patch. Given the independence of the noise, the network can only learn to determine the locally dependent true signal part of the pixel, effectively learning to denoise the volume data.

For our 3PM-Noise2Void method, we used 2% of the pixels as blind-spots, which were masked with randomly sampled pixel values from the adjacent neighborhood. A combination of L1 and L2 loss metrics was used to measure the difference between the blind-spot values in the prediction and raw volumes. We employed an ADAM optimizer^55^, with ꞵ1=0.5, ꞵ2=0.99, and a learning rate of 1e^-3^, to minimize this loss value over 300 epochs. The input volumes were randomly cut out into 16×64×64 patches, which were further augmented by rotation and flipping before applying the masking.

#### 3PM data post-processing

The 3PM data also contains periodic structured noise caused by the ripple noise of the photomultiplier tubes (PMTs), which is modulated by the line scanning process into a line-wise periodic signal. This noise violates the assumption of pixel independence and causes the 3PM-Noise2Void approach to restore or even amplify it, leading to errors in downstream analysis (see Fig. 3c). To address this problem, we optimized the hyperparameters of the 3PM-Noise2Void, identifying the z-size parameter as a key factor. We found that a depth of 16 slices minimizes the restored structured noise. However, in cases where this was not sufficient, we developed a post-processing method, referred to as PerStruc-Denoiser.

This method accounts for the origin of the noise by aligning and subsequently extracting the structured noise that occurs misaligned row-by-row and with different patterns due to the acquisition procedure and inconsistent scanning speed. The method begins by flipping each odd row of a z-slice to ensure each line is in the same scanning direction and thus has the same pattern. Next, the function determines the phase differences of the structured noise for all lines of the data to a reference line by calculating the phase shift at the main frequency component of the structured noise in the Fourier domain. This reference line is chosen as the row in the volume with the lowest standard deviation, which presents the clearest structured noise pattern and serves as an accurate reference point for the phase detection. The PerStruc-Denoiser shifts each line under periodic boundary conditions to align the periodic structured noise. By performing a median projection along the row and then along all z-slices, the algorithm extracts the pure structured noise of a single line. This line is subtracted from the modified volume, and the alignment shifts and flips are reversed. Furthermore, a Gaussian filter with a small mask size (σ = 1 pixel) is applied line by line to suppress remaining high frequency structured noise. Finally, the median value of the data is adjusted back to the original one, effectively reducing the periodic structured noise in the 3PM-Noise2Void denoised data and improving the downstream analysis of biological structures.

### Machine learning-based classification of THG signal

The THG-signal channel of 3PM images were denoised as described above. Subsequently, both the raw images as well as the denoised images were interpolated in their z-dimension with a factor of 4. The image z stacks were processed with a bilinear interpolation in this step. Semantic segmentation with ilastik was performed on the upsampled stack of raw images, using the denoised images as a prediction mask to limit the amount of false positives caused by the noise. Ilastik “Autocontext” workflow with 2 training stages was chosen for its superior performance on noisy data^56^. The THG signal was segmented into three classes of either vessel (1), myelin (2), or background (3). All color/intensity features, edge features and texture features were used up to a standard deviation of σ = 6. Additionally, 3D-features were calculated. As described previously^31^, pixels were classified using a Random Forest classifier with manually drawn labels.

To evaluate the effect of our pre-processing, an image stack was partially segmented. Labels were exported and subsequently integrated into the customized ML workflow as described, with the exception only labels from the ML training without pre-processing were used and no additional labeling was performed.

To visualize the morphological heterogeneity of myelin, vessel and background, an Uniform Manifold Approximation and Projection (UMAP) was created based on 100 features that were most distinct between groups. Those features were identified using the FindAllMarkers function in Seurat^47^.

For validation of the THG vessel signal with intravenous FITC injection, FITC images were thresholded and converted to a masked image in Fiji to obtain a segmentation for FITC signal and background. All pixels of a selected region were then analyzed and their intensity for both groups was compared.

### Polarity analysis of single glioma cells

Tumor cell polarity was investigated analyzing the primary orientation of each tumor cell in Fiji^46, 48^ and calculating the angle between the tumor cell orientation and a horizontal line. In very dense tumor regions, a subset of slices was analyzed to unequivocally identify single tumor cells. Subsequently, for each region in the corpus callosum, the overall direction of fibers was analyzed in the same manner and the calculated angles were subtracted. The angles were recalculated so that all angles were in the range of –90 to 90 degrees.

### Sample preparation, microscopy and analysis for scanning electron microscopy

Tumor-bearing PDX mice were transcardially perfused and tissue was obtained as previously described^3, 6^. For the postfixation, the mouse brain was put into 4% (w/v) PFA overnight and afterwards stored at 4°C in PBS. The brain was subsequently cut into 200 µm thick sections. To unequivocally identify the glioma cells in scanning electron microscopy, immunolabeling with an antibody against human-specific nestin (abcam, ab2035) was conducted as described previously^3^. For this purpose, the brain slices were put into 10% (w/v) sucrose dissolved in PBS for 10 minutes and for further 10 minutes in 20% (w/v) sucrose dissolved in PBS. The slice was subsequently incubated in 30% sucrose (w/v) over 12-15h at 4°C. Freeze-thaw-cycles were performed in liquid nitrogen twice for 5 minutes. Afterwards, the slices were incubated in 5% FBS in PBS at RT for 1 hour. This was followed by incubation overnight at 4°C with a human-specific mouse anti-nestin antibody (abcam, ab22035) in the blocking solution. It was washed three times with blocking solution afterwards. The slices then were incubated for 12-15h at 4°C with a secondary antibody (a biotinylated anti-mouse antibody). Slices were washed in PBS for three times and then incubated in the Vectastain ABC kit for 1h at RT. Afterwards, the samples were exposed to a solution of glucose and DAB (with glucose at a concentration of 2 mg/ml, and DAB at a concentration of 1.4 mg/ml dissolved in PBS) for 10 minutes. This was followed by an hour-long incubation in a glucose-DAB-glucose oxidase solution (with glucose oxidase at a concentration of 0.1 mg/ml, from Serva) to generate an electron-dense precipitate. The efficacy of the process was assessed using widefield light microscopy.

The labeled sections were embedded in resin and mounted onto silicon wafers as described previously^6^. For image acquisition, we used a LEO Gemini 1530 scanning electron microscope (Zeiss) combined with an ATLAS scan generator. Potential contacts of brain tumor cells and blood vessels were observed by the identification of blood vessels according to basic ultrastructural features and the DAB (diamniobenzidine)-precipitate in the brain tumor cells respectively. For three-dimensional reconstruction of blood vessel-TM contacts, images were taken at the same position in consecutive layers. A working distance of 2-4 mm was set at an aperture of 20 µm and an acceleration voltage of 2kV. The pixel sizes were between 3.8 nm and 15 nm in 400-3600 µm^2^ big images. This allowed us to create an image stack with a z-resolution of 280 nm. The manual segmentation of the structures was performed in Fiji.

### Somatokinesis measurement

Somatokinesis measurements were performed in ImageJ/Fiji^48^. First, time-lapse images of tumor cell regions were registered using custom written registration scripts. Then, tumor cells were cropped and re-registered with the 3D Drift Correction plugin in ImageJ/Fiji^57^ when necessary. The glioblastoma cell somata were labeled using a selection tool in Fiji and the center of each cell was measured. The distance between the center points of the first and last time point was calculated and divided by the duration of the experiment to obtain the somatokinetic speed of the glioblastoma cell.

### Invasion phenotype classification

Invasion phenotype classification was performed as previously described^3^. Invasion phenotypes for each cell were classified based on their invasive behavior during the time course of 4 hours.

### Tumor microtube and small process identification

A tumor microtube was defined as a small cellular protrusion with a minimum length of 10 µm and a thickness ranging between 0.5 – 2.5 µm as previously described^15^. Small processes were defined as cellular protrusions smaller than 10 µm in length as previously described^3^.

### Directionality analysis of brain tumor networks

Tumor cell regions in the corpus callosum and in the cortex were imaged with 3PM and 2PM, respectively. For each slice, the local orientation was calculated using a local window of σ = 3 pixels and a cubic spline as gradient, as implemented in OrientationJ^58^. The calculated output was exported as a stack, displaying the local orientation in degrees as well as table. For visualization of local network orientation values using rose plots, calculated orientation values were filtered 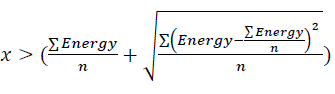 to select only tumor cell associated orientation values. For further visualization, images of tumor cell regions were binarized to generate a mask for the tumor cell network. The absolute values of the stack displaying the local orientation in degrees were multiplied with the tumor cell mask to receive the local orientation of the tumor cell network. The difference to the horizontal line was calculated for each local orientation and visually displayed as a spectrum from 90 degrees (lowest intensity) to 0 degrees (brightest intensity) in the maximum intensity mode of Arivis. This was performed for both cortex as well as corpus callosum brain regions.

### White matter reactivity analyses

Multiple regions with tumor infiltration were analyzed in the white matter-rich of the corpus callosum. Regions without THG signal in the corpus callosum were analyzed in Fiji using the polygon tool and measuring their shape descriptors on selected slices. Holes were labeled as either tumor cell-infiltrated or without tumor infiltration. Subsequently, circularity of both groups was compared.

Orientation analysis of myelinated fibers was analyzed using OrientationJ^58^. Vector fields were visualized as arrows in R.

To analyze displacement of myelin fibers by TMs or somata, TMs and somata were selected. Then, THG signal was manually labeled as displaced or non-displaced, based on THG signal. The distribution of displacing and non-displacing TMs and somata per group was analyzed.

### MRI imaging

MRI scans were conducted using a 9.4 Tesla horizontal bore small animal MRI scanner (BioSpec 94/20 USR, Bruker BioSpin GmbH, Ettlingen, Germany) equipped with a gradient strength of 675 mT/m and a 2 x 2 surface array receive-only coil. During the imaging procedure, anesthesia was induced using 4% isoflurane (Baxter, Unterschleißheim, Germany) in 100% O2, and subsequently maintained with 1-1.5% isoflurane in 100% O2 delivered via a nose cone throughout the scanning process. The respiration rate was continuously monitored, and the animals were positioned in a prone position on a Bruker standard MRI bed with an integrated circulating water heating system to ensure body temperature maintenance. A diffusion tensor imaging sequence in axial planes was employed, using the following parameters: echo time = 17.57 ms, repetition time = 1000 ms, acquisition matrix = 75×100, field-of-view = 15×20 mm2, slice thickness = 0.5 mm.

### Diffusion tensor imaging analysis

Fractional anisotropy (FA) maps were extracted from the respective DTI image stacks using open-source software MITK-Diffusion (https://github.com/MIC-DKFZ/MITK-Diffusion/)^59^. Four different regions of interest (ROI) of the corpus callosum (CC) were manually drawn by J.S. (1 year of experience in neuroimaging) and F.T.K. (board-certified neuroradiologist with 11 years of experience in neuroimaging) using the open-source software 3D Slicer (https://www.slicer.org/)^60^ at four different cross-sections of the CC moving cranially from the anterior forceps of the CC.

### 3D renderings and visualization

Renderings were created in Arivis Vision 4D for 3D and 4D Image visualization.

## Statistical analysis

The results of quantifications were transferred to GraphPad Prism (GraphPad Software) or R to test the statistical significance with the appropriate tests as indicated in the figure legends, normality was tested using the Shapiro-Wilk test. Results were considered statistically significant if the P value was below 0.05. Quantifications were done blinded by 2 independent investigators. Animal group sizes were as low as possible and empirically chosen and longitudinal measurements allowed a reduction of animal numbers by maintaining an adequate power. No statistical methods were used to predetermine sample size. In quantifications that were depicted as boxplots, the upper and the lower hinges correspond to the third and the first quartiles.

## Data availability

The source data are provided with this paper.

## Code availability

The indirect AO routines developed for integration into ScanImage32 are available at https://github.com/prevedel-lab/AO.git as previously described^22^.

AI denoising was performed using a customized approach as described above. Code can be found here: https://github.com/prevedel-lab/Deep3P-Denoising.

## Extended Data Figures

**Extended Data Fig. 1.**
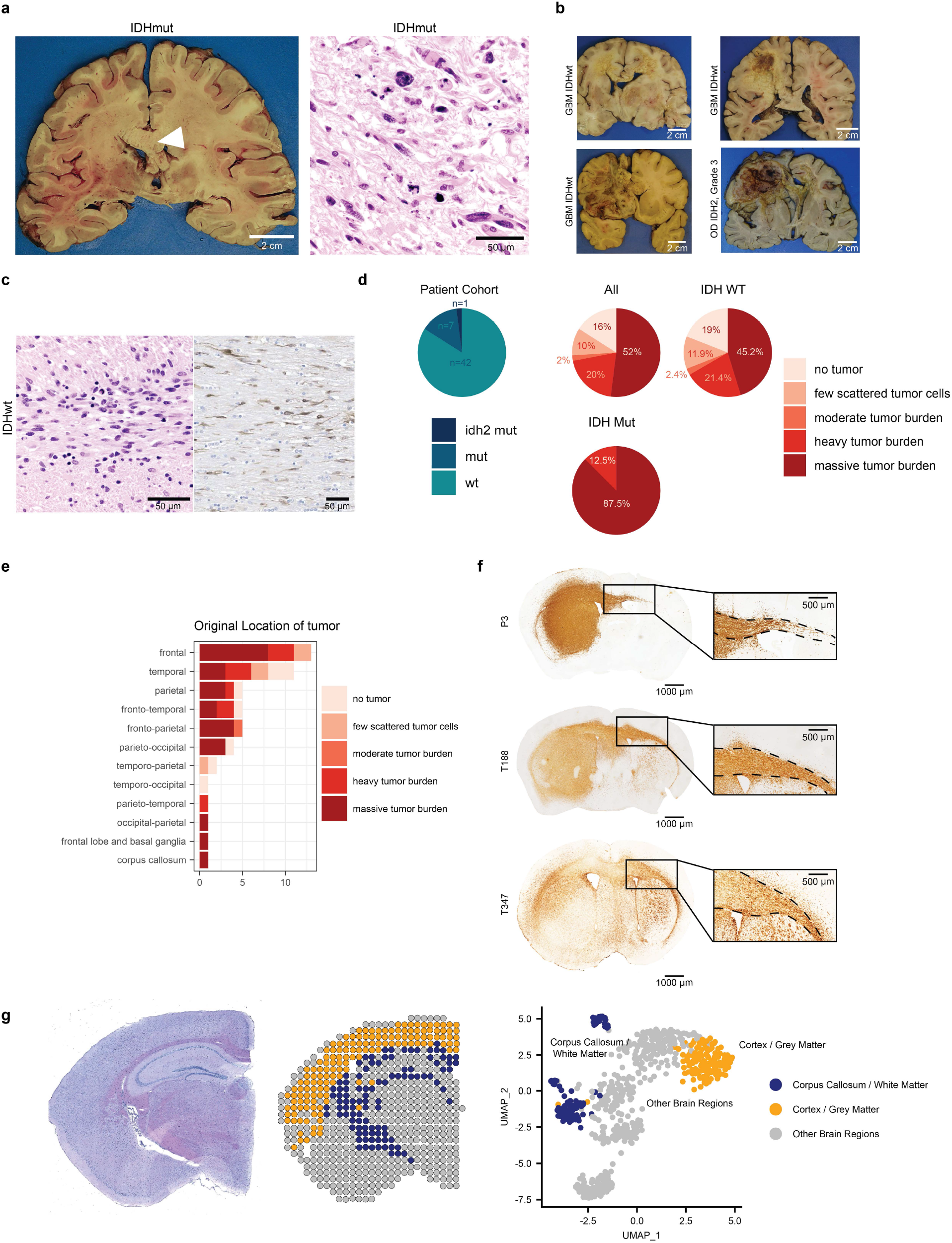
Colonization of corpus callosum as a hallmark of glioma growth patterns. a, Human autopsy from a patient with an IDH-mutant glioma and the associated histological sample. Arrowhead shows tumor infiltration in the corpus callosum. **b,** Coronal sections of brain tumor patients with IDH-WT glioblastoma and oligodendroglioma, WHO grade III showed streaks of yellow necrosis through the corpus callosum. **c**, On microscopic examination, sections of the corpus callosum showed extensive tumor infiltration on H&E (left) and in immunohistochemistry for nestin (right). On the left, histological sample from autopsy section in Fig 1a is shown. **d,** Analysis of patient cohort regarding IDH mutation (n = 50 patients). **e,** Analysis of original location of tumor and their CC infiltration. The number of patient samples is displayed on the x-axis. **f,** Examples of three different PDX models stained with an antibody against nestin and/or vimentin for the visualization of glioblastoma cells and subsequent DAB precipitation (from top to bottom: P3, T0188, T347) illustrating the infiltration into the corpus callosum. Infiltration of the corpus callosum can be seen on the inset. **g,** Spatial transcriptomic slice based on the Molecular Atlas of the Mouse Brain^46^ (left), and their molecular heterogeneity (right), visualized as UMAP (n = 639 spots).

**Extended Data Fig. 2.**
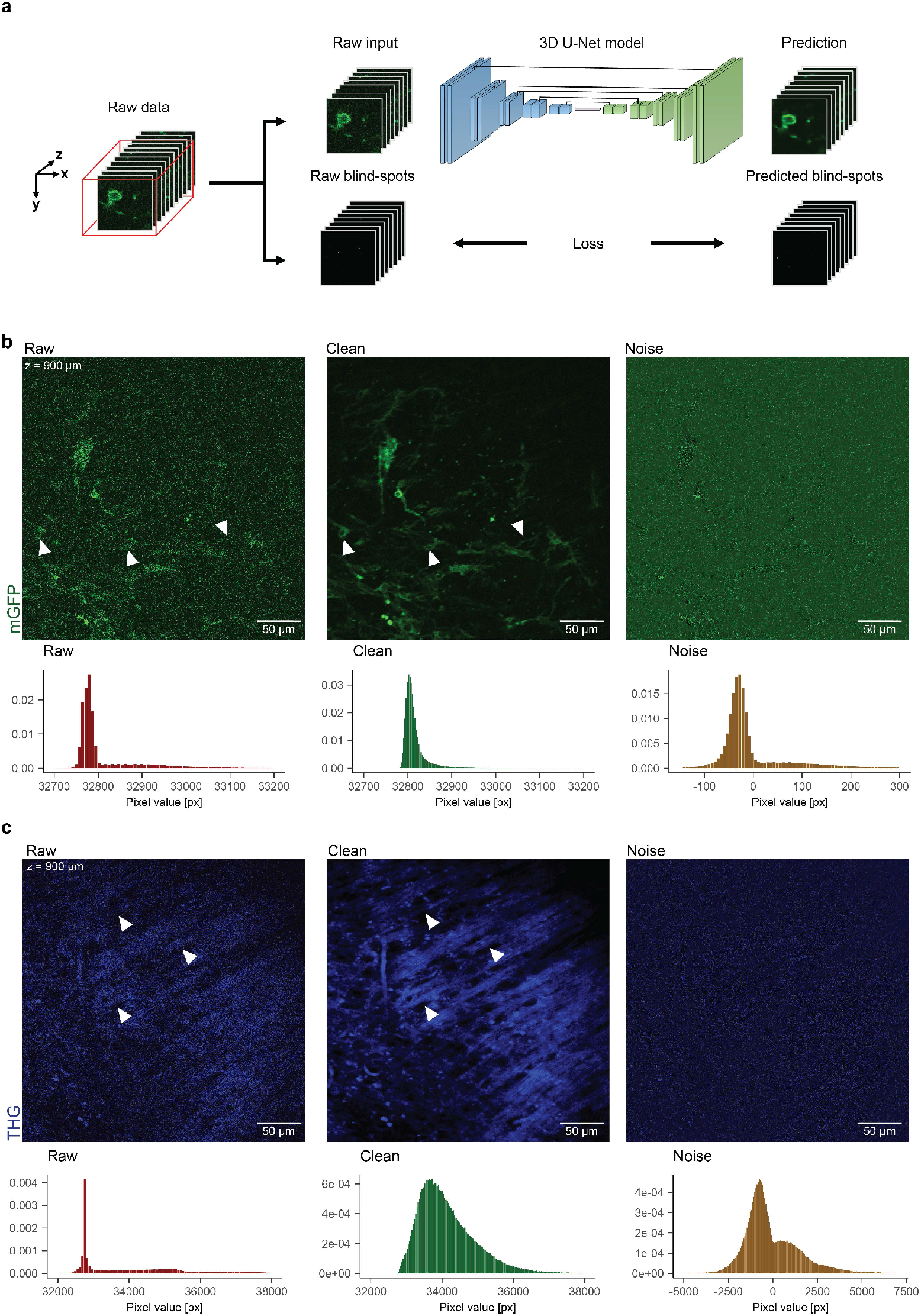
Training procedure of the three-dimensional 3PM-Noise2Void and the noise characteristics of the 3PM two-channel recording. a, Schematic representation of the 3PM-Noise2Void during training. Before the raw data is blind spotted and used for training, the three-dimensional 3PM data is sliced into smaller patches. The 3D U-net model is trained to minimize the discrepancy between the raw and predicted blind-spots. **b,** Clean and noise components of the 3PM mGFP signal with low SNR, illustrating the detector and shot noise and their narrow distribution. The clean images were estimated by averaging 60 raw images. The arrowheads point to exemplary glioblastoma cell somata and TMs. **c,** Clean and noise components of the 3PM THG signal with high SNR, showing the effects of the same noise sources and their broader distribution. The arrowheads point to exemplary myelin fibers.

**Extended Data Fig. 3.**
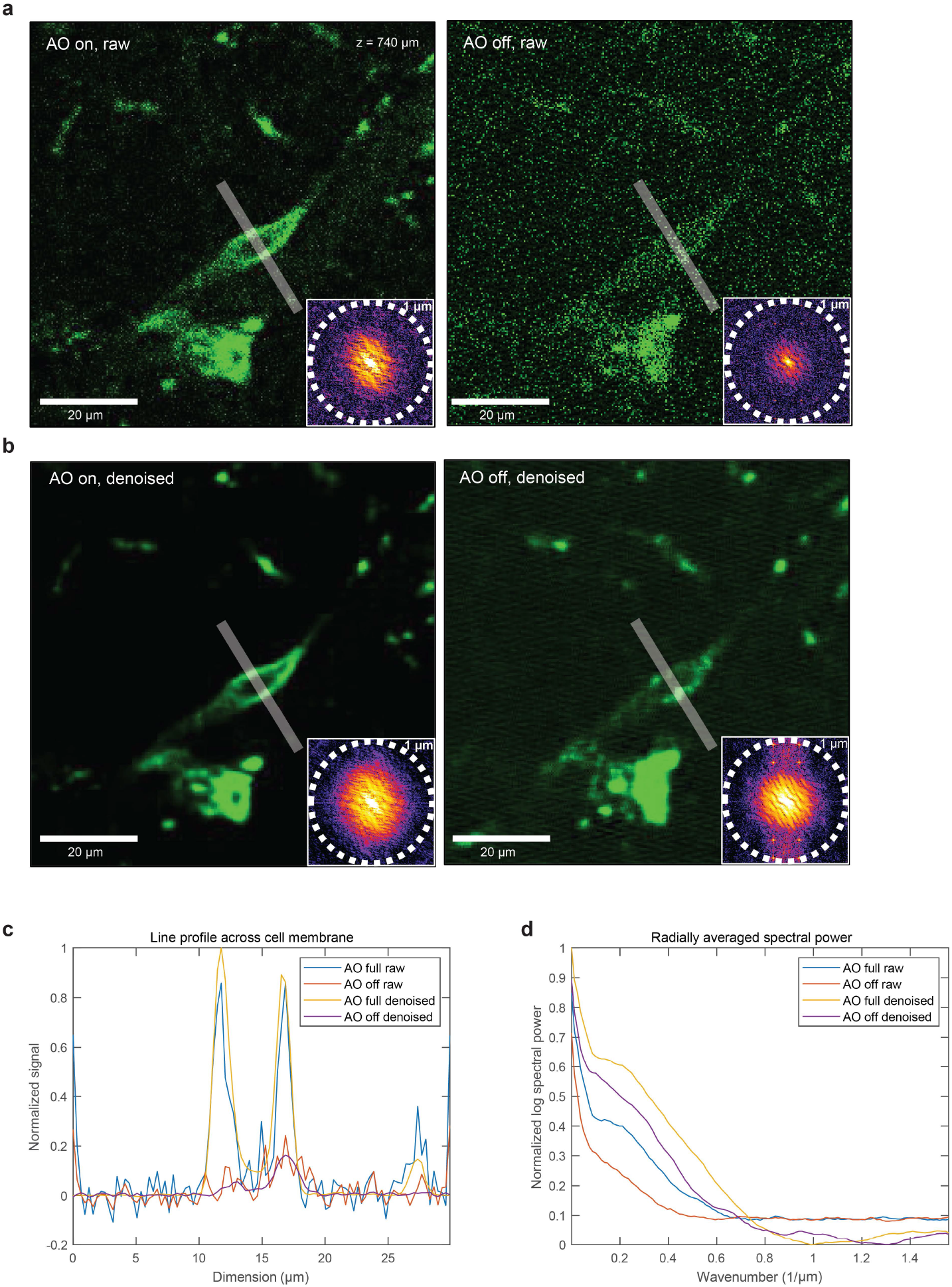
Effects of combining adaptive optics with AI-based denoising. a, An example of an mGFP labeled glioma cell imaged with (left) and without (right) AO optimization. **b,** The same panels are shown after denoising. The gray line indicates the line segment averaged over to produce the line profiles, showing the effect of uncorrected optical aberration on the visibility of fine cellular structure. The inset on each panel in **a** and **b** show the frequency domain power spectrum of the image. The image with adaptive optics shows significantly more high frequency contributions compared to the image without adaptive optics. **c,** Line profile comparisons showing intensity enhancement of the AO images. **d,** The averaged radial profile of the frequency maps are shown, allowing easier estimation of the respective frequency cut-offs.

**Extended Data Fig. 4.**
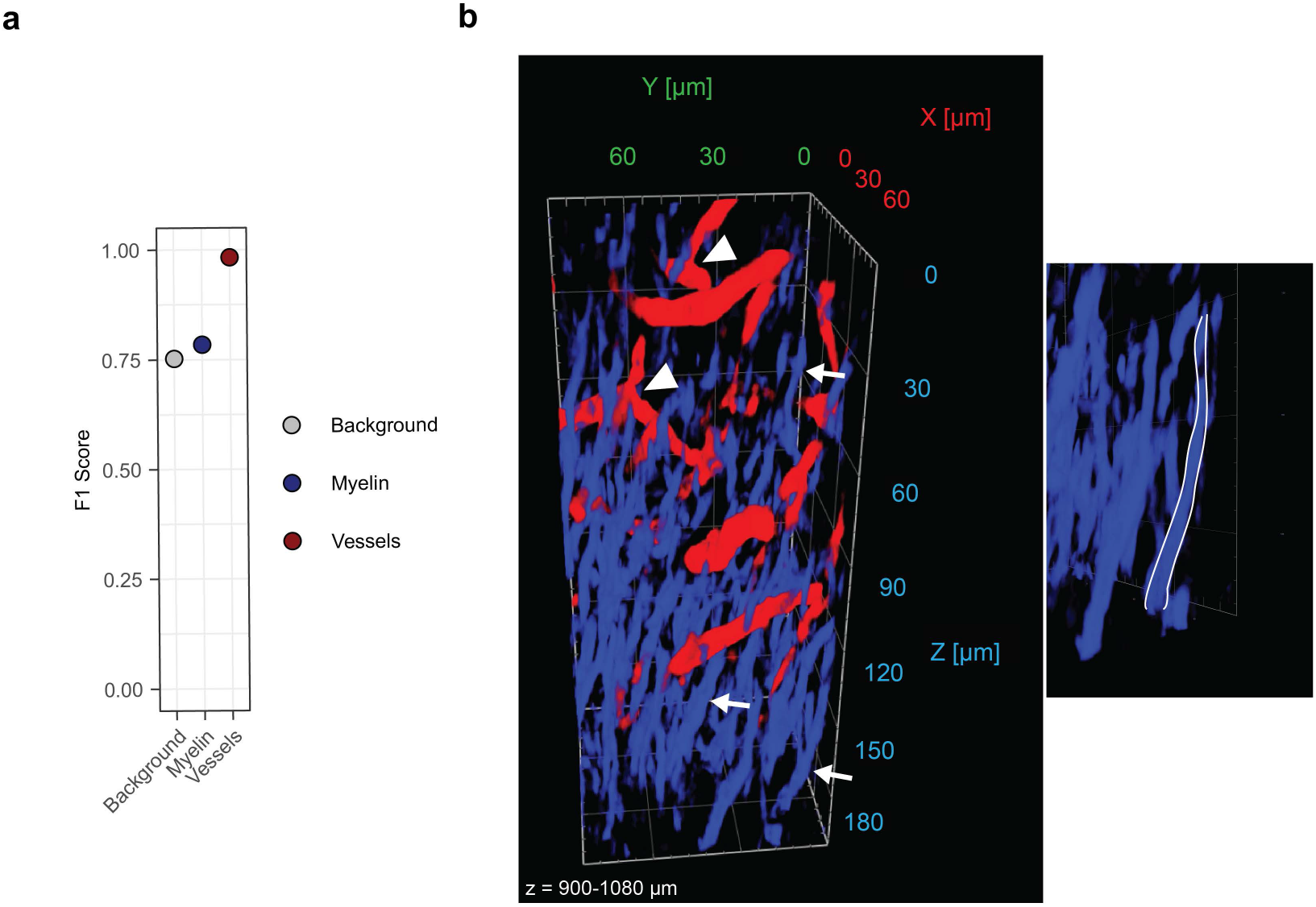
Validation of machine learning-based classification of the THG signal. a, F1 score of the machine learning based-classification as compared to the human annotated ground truth signal classification. **b,** 3D rendering of blood vessels (red) and myelinated fibers (blue) in deep cortical regions, classified using customized machine learning. Arrowheads indicate blood vessels, arrows indicate vertical myelin fibers. Myelinated fibers in the deep cortex going to the corpus callosum can be seen on the right side. Gamma values were adjusted for 3D visualization.

**Extended Data Fig. 5.**
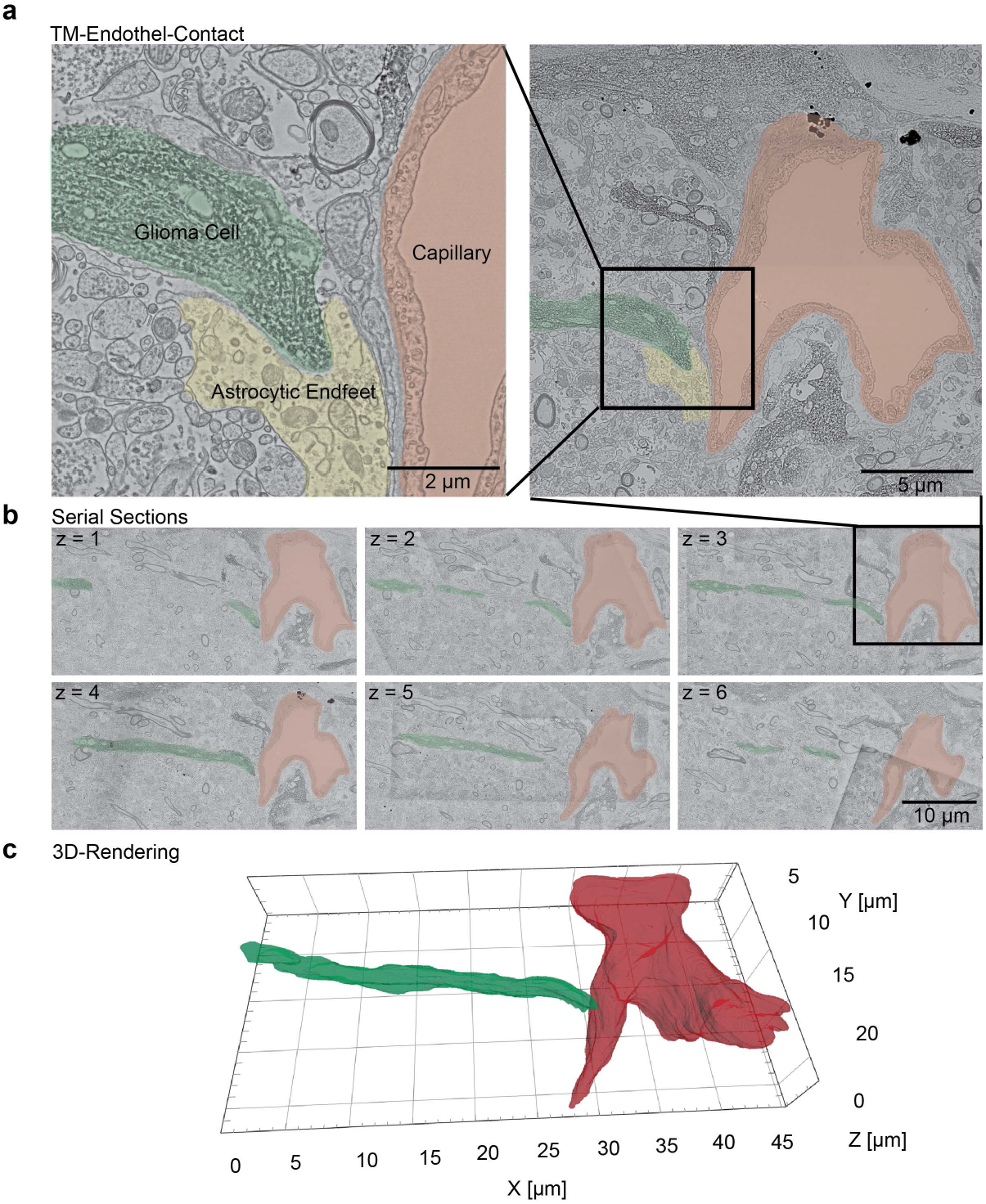
3D electron microscopy reconstruction of vessel-guided glioblastoma translocation. a, Scanning electron microscopy image of an orthogonal contact of a TM (green) on a blood vessel (red). Astrocytic endfeet colored in yellow, cell type based on ultrastructural features. Zoom-in on the left side (n = 13 GBCs from n = 5 PDX mice). DAB-precipitate at anti-nestin-stainings. **b**, Serial section of scanning electron microscopy images showing the GBC-vessel-contact. **c**, 3D rendering of the TM-tip on the capillary using manual segmentation of 20 serial 2D-EM-sections. Gamma values were adjusted for 3D visualization.

**Extended Data Fig. 6.**
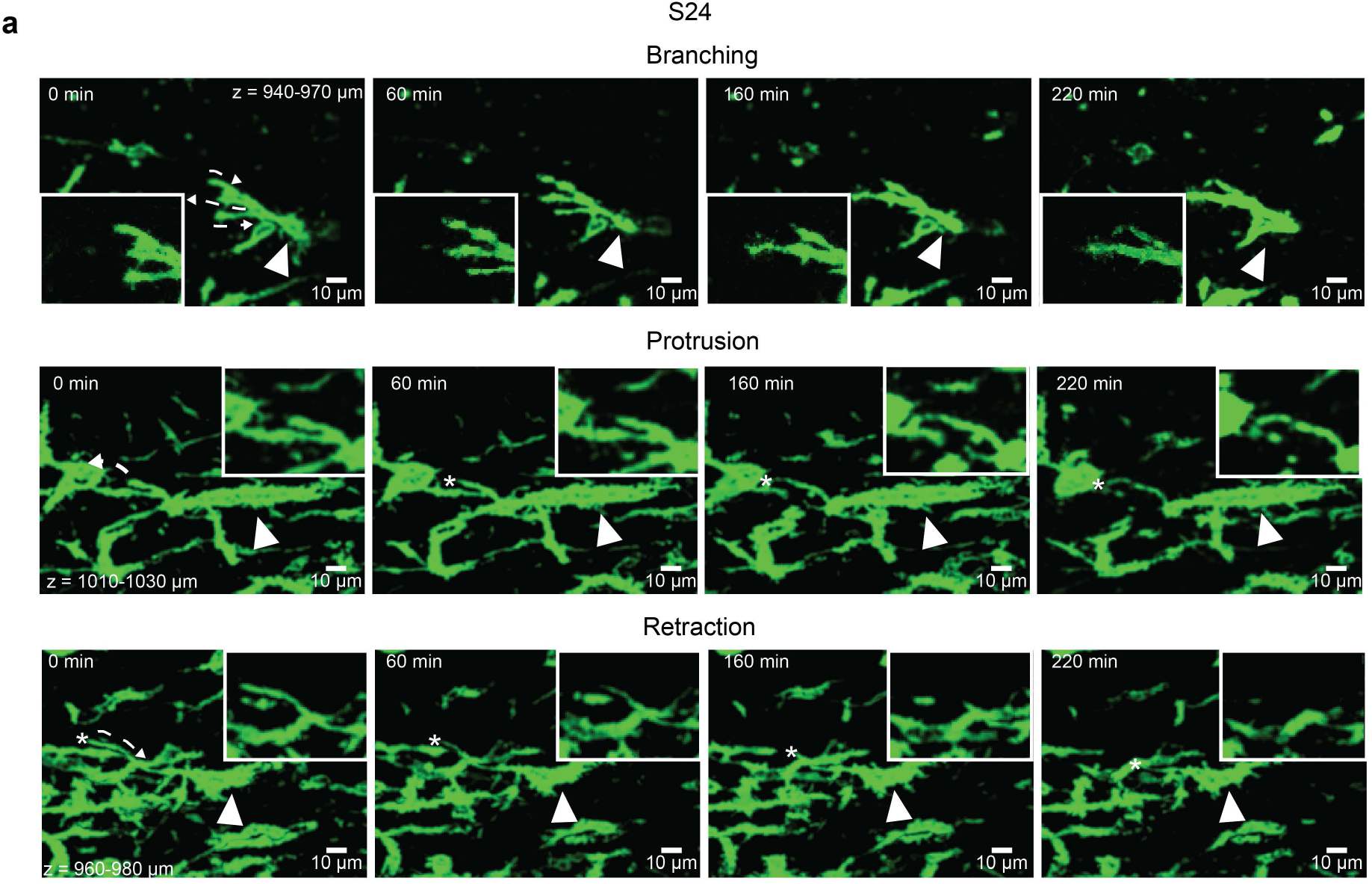
Tumor microtube dynamics in the corpus callosum in further PDX models. a, Maximum intensity projection time-lapse images showing TMs that use branching, protrusion, and retraction in the corpus callosum in the PDX model S24 (n = 3 S24 PDX). Arrowheads point at the somata, the dashed arrows indicate the direction of the tumor microtube dynamic. Data are shown as probability maps and post-processed with the ‘‘smooth” function in Fiji.

**Extended Data Fig. 7.**
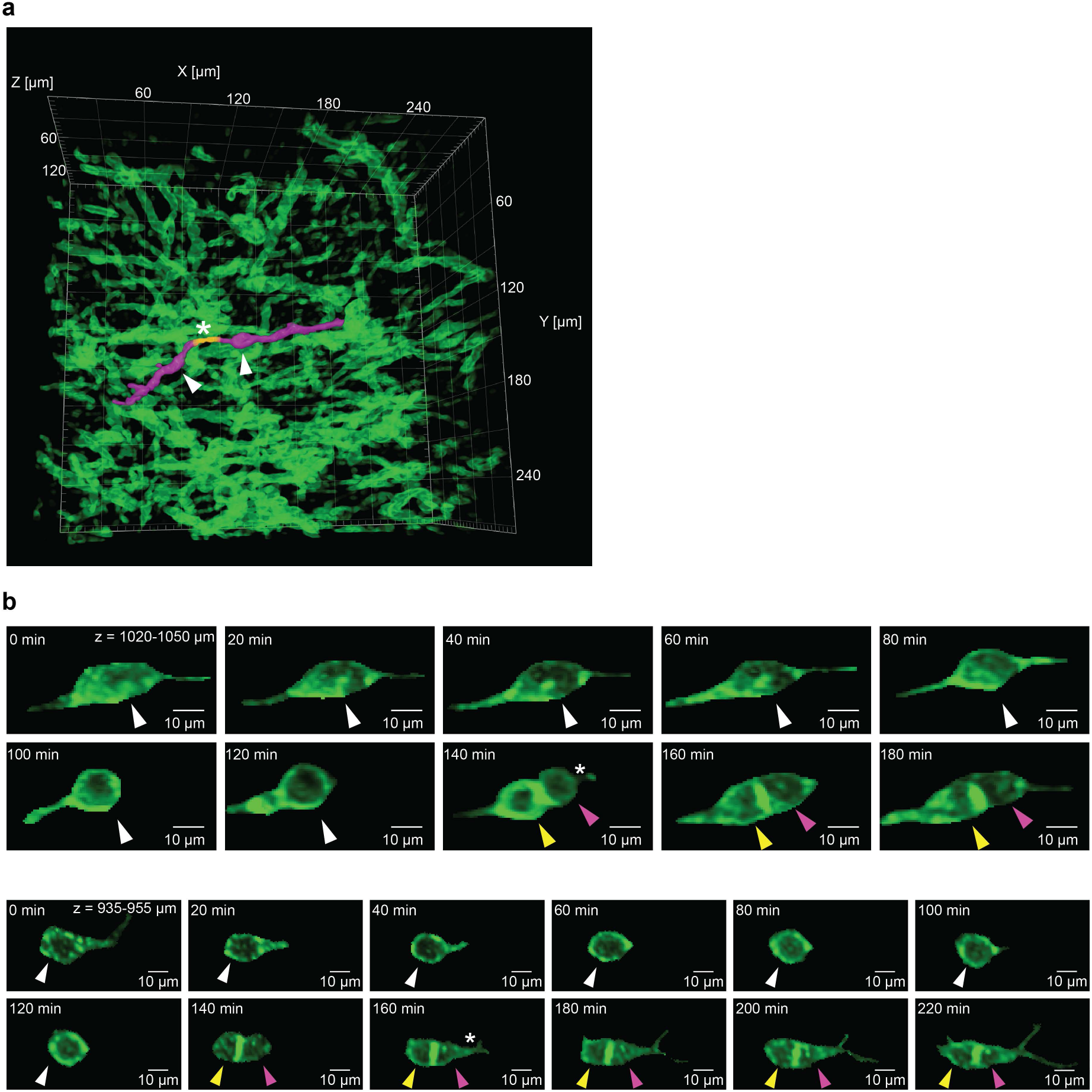
Time-lapse of tumor cell proliferation in the corpus callosum. a, Rendering of tumor cell network. An exemplary tumor-tumor cell connection is visualized (yellow, asterisk) that connects two tumor cells (purple, arrowhead). Gamma values were adjusted for 3D visualization. **b,** Maximum intensity time-lapse of tumor cell proliferation in the corpus callosum. White arrowhead: Glioblastoma cell before cell division. Yellow and purple arrowhead: Daughter glioblastoma cells after cell division. The asterisk points at a newly grown TM. Top: S24 PDX model, bottom: T269 PDX model. Post-processed with denoising and “clear outside” function in ImageJ/Fiji.

**Extended Data Fig. 8.**
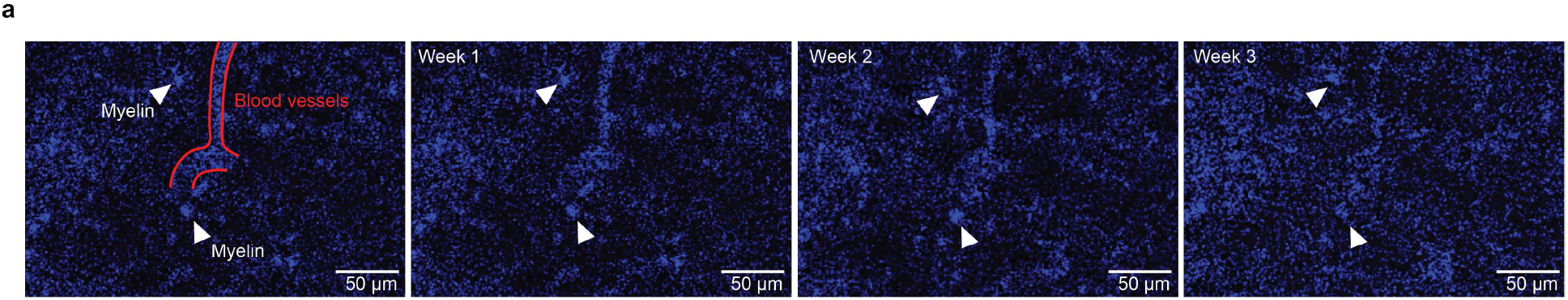
Following myelin tracts and blood vessels over weeks in 3P. a, Arrow heads indicate myelin fibers, blood vessel is labeled in red (left). Post-processed with denoising.

**Supplementary Table 1:** Data of glioma patients analyzed for macroscopic and microscopic infiltration of the corpus callosum.

**Supplementary Table 2:** Molecular characterization of patient-derived glioblastoma cell lines and PDOX models used in this study.

**Supplementary Table 3:** Acquisition parameters for high resolution *in vivo* imaging at different depth shown in this work.

**Supplementary Video 1:** A three-dimensional rendering of a patient-derived glioblastoma xenograft model (S24) in cortex and corpus callosum with 3PM. Visualization of glioblastoma cells (green), blood vessels (red) and the corpus callosum (blue).

**Supplementary Video 2:** A three-dimensional rendering of the THG signal in deep cortex layers. The signal was classified with customized machine learning into vessel (red) and myelin fibers (blue). 3PM was performed in the patient-derived xenograft model S24. Imaging depth was between 400-700 µm.

**Supplementary Video 3:** 3D rendering of near-diffraction limited imaging of glioblastoma, blood vessels and white matter tracts in the corpus callosum. Small processes of the glioblastoma cell can be seen. 3PM was performed in the patient-derived xenograft model T269. Imaging depth was between 840-920 µm.

**Supplementary Video 4:** Three neuronal-like cellular mechanisms of brain tumor invasion in the corpus callosum. 4D time-lapse imaging examples of branching migration, locomotion and translocation with Deep3P. Imaging depth was at 990-1020 µm.

**Supplementary Video 5:** Tumor-tumor networks in white matter tracts. The S24 PDX model is shown. Glioblastoma cells are depicted in green, white matter tracts in blue and blood vessel signal in red. Imaging depth was between 900-1060 µm.

**Supplementary Video 6:** Glioblastoma cell division in the corpus callosum, followed over 180 minutes. The S24 PDX model is shown. Cell is colored green before division, two daughter cells after division are colored in yellow and purple. Imaging depth was at 985-1010 µm.

<u>Supplementary Note 1:</u> Deep3P imaging considerations and workflow

Due to the limits on pulse energy and average power that can be safely delivered^51^ to the focal volume during imaging, it is important to have optimized window placement and properties as well as a refined acquisition plan in order to find and characterize volumes of interest. In case that no labeled cells are found initially, it is not clear whether they are absent, or the signal to noise is simply too poor to resolve them. Conducting the exploration while exceeding recommended laser power levels may result in damage to the brain regions explored as well as bleaching the fluorescent label in those volumes. Having an additional structure labeled, or in the case of this work, a label-free signal like THG, greatly aids in ensuring that absence of labeled cells is not mistaken for poor signal to noise.

Poor signal to noise performance can result from microscope issues (uncompressed pulses, clipped or distorted beam profiles, misalignment etc.), window issues (optical quality, curvature, glue/dirt obscuration, inflammation, bad placement etc.), or from brain structures obscuring the infiltrated areas (large blood vessels or other absorbing structures). All three sources can also produce a distorted PSF due to strong aberration, resulting in low SNR for 3PM, and which in extreme cases cannot be fully mitigated by AO optimization.

It is therefore critical that the cranial window in of the highest optical quality, placed over a region of the brain ensuring wide and deep access to the most accessible white matter structures, and that the glue is both secure and not obscuring or contaminating the imaging area. The longitudinal 2PM imaging of glioblastoma in the mouse brain is helpful in tracking the progress of tumor progression. Subsequently, the goal is then to optimize access to as much white matter as possible that yields desirable SNR properties. The inhomogeneous illumination in cortex and the corpus callosum also means that simply turning up the laser power is not advisable. One hemisphere can be privileged for wider or deeper access by moving the window off-center from the sinus midline towards the target hemisphere. Therefore, an imaging experiment proceeds with identifying the accessible patches, then acquiring volumes to screen for labeled cell infiltration. This proceeds most easily by locating the depth of the window surface, the lateral position of the sinus, and then moving 1200 µm from the sinus laterally to a relatively unobstructed cone of light. Proceeding to roughly 900 µm depth (ensure the objective correction collar is set to this depth), the power can be increased gradually to 80% or so of the threshold power expected for that depth and brain anatomy, as estimated by taking the effective attenuation length of brain tissue into account. As the power is brought up and travels lengthwise with the objective, THG signal from the white matter will increase dramatically when an accessible patch is encountered. Once a path is found, higher resolution characterization reveals the depth and thickness of the CC, identified by visible fiber alignment. Once the desired imaging volume is thus identified, labeled cells can either be directly screened, or an AO optimization on either the white matter THG or on any labeled cells can be performed. Optimizing the AO before intensive characterization helps ensure that the minimum necessary power is used, minimizing light exposure and enabling a four hour time-lapse experiment to be conducted without photodamage or significant bleaching.

